# ERα-LBD, a novel isoform of estrogen receptor alpha, promotes breast cancer proliferation and endocrine resistance

**DOI:** 10.1101/2021.10.14.464262

**Authors:** Antonio Strillacci, Pasquale Sansone, Vinagolu K Rajasekhar, Mesruh Turkekul, Vitaly Boyko, Fanli Meng, Brian Houck-Loomis, David Brown, Michael F Berger, Ronald C Hendrickson, Qing Chang, Elisa de Stanchina, Fresia Pareja, Jorge S Reis-Filho, Ramya Segu Rajappachetty, Bo Liu, Alex Penson, Chiara Mastroleo, Marjan Berishaj, Francesca Borsetti, Enzo Spisni, David Lyden, Sarat Chandarlapaty, Jacqueline Bromberg

## Abstract

Estrogen receptor alpha (ERα) drives mammary gland development and breast cancer (BC) growth through an evolutionarily conserved linkage of DNA binding and hormone activation functions. Therapeutic targeting of the hormone binding pocket is a widely utilized and successful strategy for breast cancer prevention and treatment. However, resistance to this endocrine therapy is frequently encountered and may occur through bypass or reactivation of ER-regulated transcriptional programs. We now identify the induction of a novel ERα isoform, ERα-LBD, that is encoded by an alternative *ESR1* transcript and lacks the activation function and DNA binding domains. Despite lacking the transcriptional activity, ERα-LBD is found to promote breast cancer growth and resistance to the ERα antagonist fulvestrant. ERα-LBD is predominantly localized to the cytoplasm and mitochondria of BC cells and leads to enhanced glycolysis, respiration and stem-like features. Intriguingly, ERα-LBD expression and function does not appear to be restricted to cancers that express full length ERα but also promotes growth of triple negative breast cancers and ERα-LBD transcript (ESR1-LBD) is also present in BC samples from both ERα(+) and ERα(−) human tumors. These findings point to ERα-LBD as a potential mediator of breast cancer progression and therapy resistance.

**SIGNIFICANCE STATEMENT:** Endocrine resistant and metastatic breast cancer (BC) is a clinically significant problem. Our study of fulvestrant resistant cancer cells led to the discovery of a novel ERα isoform which we call ERα-LBD. Encoded by a truncated transcript variant (ESR1-LBD) and lacking the N-terminal domains (activation of transcription and DNA binding), ERα-LBD displays a unique role in BC tumorigenesis and progression by mechanisms that may involve metabolic and cell growth advantages, stemness and therapy resistance. Importantly, ESR1-LBD is preferentially expressed in human breast tumor tissues and may be used as prognostic marker in BC.

## INTRODUCTION

The nuclear receptor estrogen receptor alpha (ERα) is the major therapeutic target for the ~75% of breast cancers (BC) where its expression is detected (**1**). ERα is a multifunctional protein that contributes to cellular processes via transcriptional regulation, participation in signaling complexes (**2**, **3**) and regulation of mitochondrial function (**4**, **5**). Its function in primary BC growth depends mainly on its activation of transcription in response to hormone binding-mediated conformational changes in the receptor. Thus, therapeutic targeting of ERα has been directed at inhibiting this hormone-receptor interaction. The major forms of therapy include suppressors of estrogen biosynthesis (*e.g.* aromatase inhibitors, GNRH antagonists) and direct ERα antagonists (*e.g.* tamoxifen, fulvestrant) (**6**–**7**). Cancers develop resistance to these forms of therapy over time, often through alterations in the *ESR1* gene including activating point mutations in the ligand binding domain and gene fusions (**8**–**10**) that restore ERα transcriptional activity. In addition, cancers can bypass ERα signaling and activate oncogenic functions through genetic alterations that involve other growth factor signaling pathways, modification of cell cycle regulators and increases in stem cell activity (**11**–**14**). Additionally, epigenetic mechanisms of resistance may also be contributory and whether these can affect ERα or these other growth pathways is a key question, particularly given that up to half of all resistant tumors harbor no mutation known to cause resistance (**10**, **15**).

About 15% of BC cases are triple-negative breast cancers (TNBCs), lacking the expression of ERα, PR and HER2-amplification and are thus not responsive to targeted therapies against ERα or HER2. These cancers are associated with a poorer prognosis due to a higher rate of metastatic progression and resistance to treatment (**16**, **17**). Notably, TNBC may be responsive to estrogen stimulation via ERα-independent pathways, promoting tumor formation and progression via different molecular mechanisms (**18**).

In this work, we investigated ERα protein and transcripts in *ESR1* wildtype breast cancers that were resistant to fulvestrant and in TNBC. We identified an isoform that maintains the ligand binding domain but lacks the DNA binding domain, and promotes breast cancer growth and endocrine resistance through non-canonical functions outside the nucleus. These data reinforce the importance of the broader biological functions of ER protein family outside of its transcriptional activation role.

## RESULTS

### Fulvestrant resistant cells express a truncated estrogen receptor α isoform

In order to investigate novel mechanisms of hormonal therapy resistance (HTR) in breast cancer, we examined the expression of ERα protein in different BC cell lines, either in the absence or presence of the selective estrogen receptor degrader (SERD) fulvestrant (ICI 182,780) which reduces ERα full-length (ERα-FL, 66 kDa) protein levels as a consequence of reduced stability and dimerization (**19**). We postulated that changes in fulvestrant-mediated suppression of ERα-FL levels may promote drug resistance and investigated this response in several models selected for fulvestrant resistance (MCF-7 FulvRes, MCF-7 Y537S and two PDX models) (**14**, **20**), compared to fulvestrant-sensitive MCF-7 cells (**Fig. 1A**; **Supplementary Fig. S1A** and **B**). After 24 hours exposure to 1 mM fulvestrant, MCF-7 cells and MCF-7 FulvRes showed ~80% and ~60% loss of ERα-FL expression, respectively; MCF-7 Y537S and PDX-ERα(+) showed a slight decrease (~20% and ~30%) and PDX-ERα(−) showed no expression at all. Notably, we observed a faster migrating band (~37 kDa) in all four resistant models that appeared to increase in response to fulvestrant. As this protein was detected in both ERα positive and negative models, we analyzed by immunoblotting two triple-negative breast cancer (TNBC) cell lines, MDA-MB-453 and MDA-MB-231, also lacking ERα-FL expression and therefore resistant to fulvestrant treatment (**Supplementary Fig. S1A**). Notably, both TNBC cell lines displayed this lower molecular weight protein whose expression increased after fulvestrant treatment (**Fig. 1B**). To establish the identity of the novel protein, we performed immunoprecipitation and mass spectrometry experiments from MCF-7 FulvRes and MDA-MB-231 cells and determined that the ERα-related protein is a truncated isoform of ERα-FL, lacking the N-terminal domains AF1 (transcription Activation Function-1), DBD (DNA Binding Domain) and a portion of the hinge domain, and it is composed principally by the C-terminal domains LBD (Ligand Binding Domain) and AF2 (**Fig. 1C**; **Supplementary Fig. S1C**). Based on our observation, we named this novel ERα protein variant ‘ERα-LBD’. ERα-LBD has 332 amino acids, a MW of 37.3 kDa and a predicted 3-D structure that is similar to the predicted ERα-FL (**Fig. 1C; Supplementary Fig. S1D**). As expected, we failed to detect a 37.3 kDa protein using an ERα antibody raised against the amino-terminus (**Supplementary Fig. S1E**).

**Figure 1.**
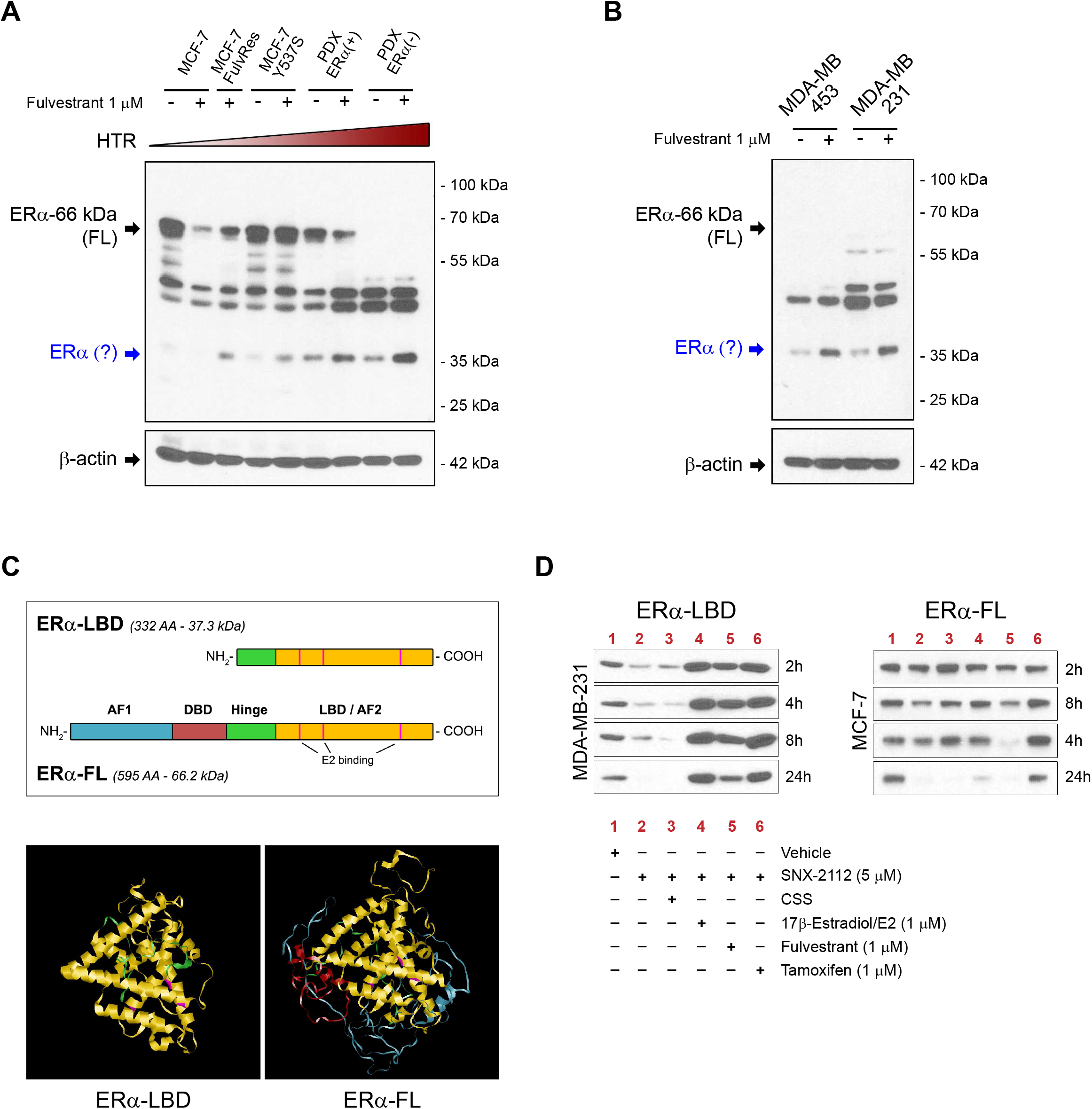
Fulvestrant resistant BC cell lines express ERα-LBD, a novel estrogen receptor α variant. **A** and **B,** Western blot analysis of estrogen receptor alpha (ERα) protein expression in breast cancer (BC) cell lines. Cells were cultured for 24 h in the presence of vehicle (−) or fulvestrant 1 μM (+) before lysis. Vehicle = DMSO. ERα protein expression was normalized against J3-actin. Increasing levels of hormonal therapy resistance (HTR) are indicated. A potential smaller ERα isoform is indicated in blue. **C,** Comparison between ERα-LBD and ERα-FL (full-length): domains (AF-1, transcription Activation Function-1; 080, DNA Binding Domain; Hinge; LBD, Ligand Binding Domain; AF-2, transcription Activation Function-2), amino acid (AA) number, molecular weight, 3-0 structure prediction (Phyre2). **D,** ERα protein stability assay. ERα-LBD and ERα-FL protein levels were analyzed by western blot in MDA-MB-231 and MCF-7 cells, respectively. Protein samples were collected at different time points (2-24 h), after treatment. Vehicle = DMSO; CSS = charcoal stripped serum.

Across all the models we investigated, we found that fulvestrant increased expression of ERα-LBD and in some cases was necessary for detection. These data suggested that fulvestrant was serving to stabilize ERα-LBD akin to the way ligands protect full-length ER against proteolytic degradation when unchaperoned by HSP90. To further address this possibility, we analyzed ERα-LBD and ERα-FL expression upon HSP90 inhibition using an ATP inhibitor of HSP90 function (SNX-2112) (**21**). Following SNX-2112 treatment, both ERα-FL and ERα-LBD underwent rapid degradation, either in the presence (lane 2) or absence (lane 3, CSS) of physiological E2 levels. Conversely, high levels of E2, fulvestrant or tamoxifen treatment (1 μM) led to increased stability of ERα-LBD. (**Fig. 1D; Supplementary Fig. S1F**).

Taken together, our data demonstrate that ERα(+) and (−) BC models characterized by HTR can produce a truncated ERα isoform whose expression can be induced by ERα ligand binding pocket compounds, including fulvestrant.

### ERα-LBD is encoded by an ESR1 transcript variant

To understand the basis of ERα-LBD formation we carried out RNA capture-sequencing and PCR assays to analyze the usage of ESR1 exons and transcripts in different models, including both ERα-FL positive and negative cells. High-depth capture-sequencing data showed that BC ERα-FL(−) cells expressing exclusively ERα-LBD, such as MDA-MB-453/-231 and PDX-ERα, have higher abundance of ESR1 read counts in the C-terminus (CDS exons #4, 5, 6, 7 and 8), compared to the N-terminus (CDS exons #1-3). In contrast, ESR1 exon usage profile was significantly different in ERα-FL(+) cells, such as T-47D, MCF-7 and PDX-ERα(+), where the expression of exons #1-3 was higher than exons #4-8 (**Fig. 2A**; **Supplementary File S1**). To identify an ESR1 transcript variant for ERα-LBD, we searched the FANTOM CAT (CAGE Associated Transcriptome) database, which displayed an additional ESR1 exon between exons #3 and #4, with a transcription start site (TSS) at the 5’ end (**Supplementary Fig. S2A**) (**22**). We annotated this region as 5’UTR of a putative ESR1-LBD transcript variant (exon E3a). In this region, a higher number of read counts was found in ERα(−) cells, compared to ERα(+) (**Fig. 2A**). Differential exon usage was determined for ESR1-FL (encoding ERα-FL) and a predicted ESR1-LBD (encoding ERα-LBD) transcript by comparing ERα-FL(+) and ERα(−) groups, revealing a significantly lower usage of exons E1, E2, E3 and E8-3UTR, and higher usage of exons E3a, E5, E6 and E7 for the ERα(−) group (**Fig. 2B**).

**Figure 2.**
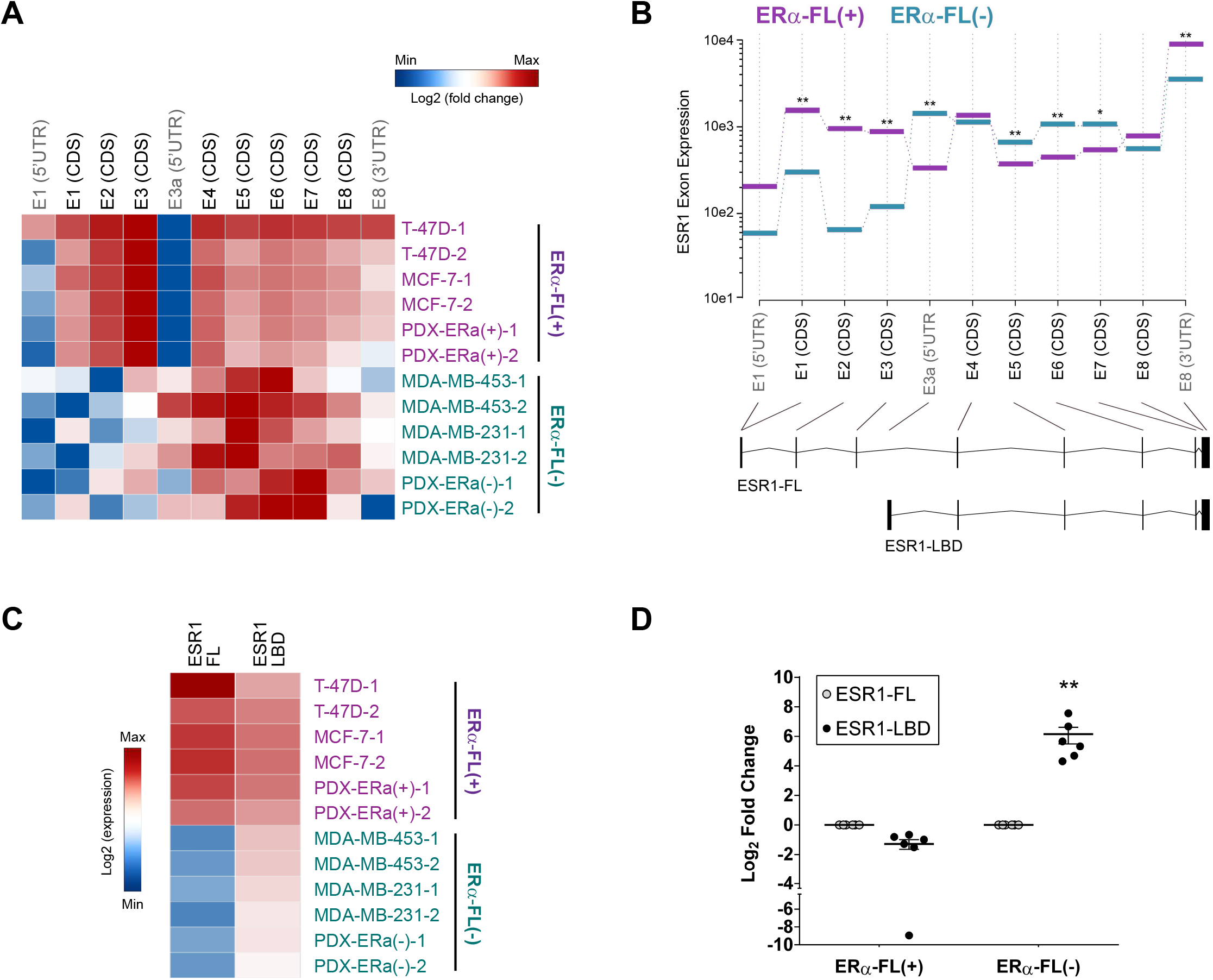
ERα-LBD is encoded by an ESR1 transcript variant. **A-D,** Total RNA was extracted from different BC cell lines and analyzed by RNA capture sequencing (capture-seq). Data output was used to investigate differential expression (DE) of ESR1 exons and transcript variants among samples. **A,** Heat map showing ESR1 exon expression calculated by using DEXSeq-count tool (log2, mean-centered). Exons encoding ERα-FL protein are indicated in black (CDS). BC cell lines were ranked based on their known ERα-FL protein expression and grouped as ERα-FL(+), including T-47D, MCF-7, PDX-ERα(+) (in purple), and ERα-FL(−), including MDA-MB-453, MDA-MB-231, PDX-ERα(−) (in teal). **B,** Differential ESR1 exon expression between ERα-FL(+) and ERα­ FL(−) groups was calculated using DEXSeq and plotted using plotDEXSeq tools. Exons associated with two different ESR1 transcript variants were taken under consideration for the analysis: ESR1-FL (main variant encoding ERα-FL) and ESR1-LBD (predicted, encoding ERα-LBD). Expression values for ERα-FL(+) and ERα-FL(−) groups are plotted in purple and in teal, respectively. BH adjusted P-values: *\*P* < 0.05, *\*\*P* < 0.01 *(n* = 6/group). **C,** Heat map showing ESR1 transcript (ESR1-FL and ESR1-LBD) expression among BC cell lines, ranked as ERα-FL(+) (purple) and ERα-FL(−) (teal). Transcript expression was calculated by using Kallisto tool and quantified as counts / transcript length. **D,** ESR1-LBD transcript expression plotted as log2 fold change, relative to ESR1-FL. Color code: grey, ESR1-FL; black, ESR1-LBD. Mean± s.e.m. is shown *(n* = 6); ** *P* < 0.01, ratio paired t test, two-sided.

Exon data were supported by the analysis of ESR1 transcripts expression, demonstrating higher levels of ESR1-LBD transcript in ERα-FL(−) cells, compared to ESR1-FL (**Fig. 2C** and **D**; **Supplementary File S1**). Also qPCR (exon ‘walking’) and RT-PCR assays confirmed the RNA-seq results. ERα-FL(+) and ERα-FL(−) cells express an ESR1 mRNA ranging from exon 1 to 8 and from exon 4 to exon 8, respectively (**Supplementary Fig. S2B** and **C**). These results are in accordance with the western blot data in **Fig. 1B**, showing that ERα-FL(−) cells express a truncated ERα protein (LBD) and not full-length ERα. As definitive proof, the stable cloning of the putative ERα-LBD CDS into BC cells (exon 4 to exon 8), led to ERα-LBD overexpression (oe) whereas targeted genomic deletions within exon 4 (by CRISPR technology) induced ERα-LBD knockdown (kd) (**Supplementary Fig. S2D**).

RNA sequencing and qPCR data showed reduced expression of the ESR1 3’UTR sequence in ERα-FL(−) cells as compared to ERα-FL(+) ones (**Fig. 2A** and **B**; **Supplementary Fig. S2B**). It has been observed that shortening of 3’UTRs by alternative cleavage and polyadenylation is associated with increased RNA stability, and that this phenomenon can promote oncogene mediated transformation by enhanced protein production (**23**). Thus, we hypothesized that the partial/total loss of the 3’UTR sequence in ERα-FL(−) cells may enhance ESR1 mRNA stability. To test this, we treated BC cells with actinomycin-D (inhibitor of RNA synthesis) and indeed the LBD portion of ESR1 transcript was significantly more stable than the FL-specific in MCF-7 FulvRes and TNBC cells (**Supplementary Fig. S2E**).

We note that ERα-LBD is not the only ERα transcript variant that has been observed in breast cancer. An ERα-36 variant, encoded by ESR1 exons #2-6 and an additional exon downstream of the ESR1 gene, encoding a protein expressing the DNA-binding domain, a truncated LBD and a partial dimerization domain has been described in a variety of cancer models including tamoxifen resistant breast cancers (**24**, **25**). Importantly, ERα-LBD isoform is distinct from ERα-36 as specific primers and an antibody for the ERα-36 variant failed to detect ERα-LBD (**Supplementary Fig. S2F** and **G**).

### ERα-LBD localizes to the mitochondria and cytoplasm

ERα-LBD lacks the N-terminal domain of the full-length receptor and we thus hypothesized it may have a distinctive cellular location. Using ERα immunohistochemistry (IHC), full-length expressing MCF-7 cells show robust nuclear staining that is markedly reduced 24 h after fulvestrant treatment. By contrast, ERα-LBD expressing models MCF-7 FulvRes, MDA-MB-453/-231 and PDX-ERα(−) display punctate cytoplasmic foci (**Fig. 3A**). Based on this particular pattern, we hypothesized that ERα-LBD might be localized to the mitochondria and performed confocal analyses examining cells for ERα protein (green), OXPHOS (mitochondrial marker, red) and DAPI (nuclear marker, blue) (**Fig. 3B**). We examined images from both separate and merged channels, and by 3-D image processing to evaluate the co-localization volume between ERα, nuclei or mitochondria. The overall IF staining confirmed data from the IHC: MCF-7 cells have a strong ERα nuclear localization which is decreased after fulvestrant treatment; MCF-7 FulvRes chronically treated with fulvestrant have both nuclear and mitochondrial ERα staining. In full-length negative cells, ERα-LBD was largely absent from the nucleus and found in the cytoplasm with a substantial fraction co-localized with mitochondria. Detailed co-localization values and statistics are shown in **Supplementary Fig. S3A**.

**Figure 3.**
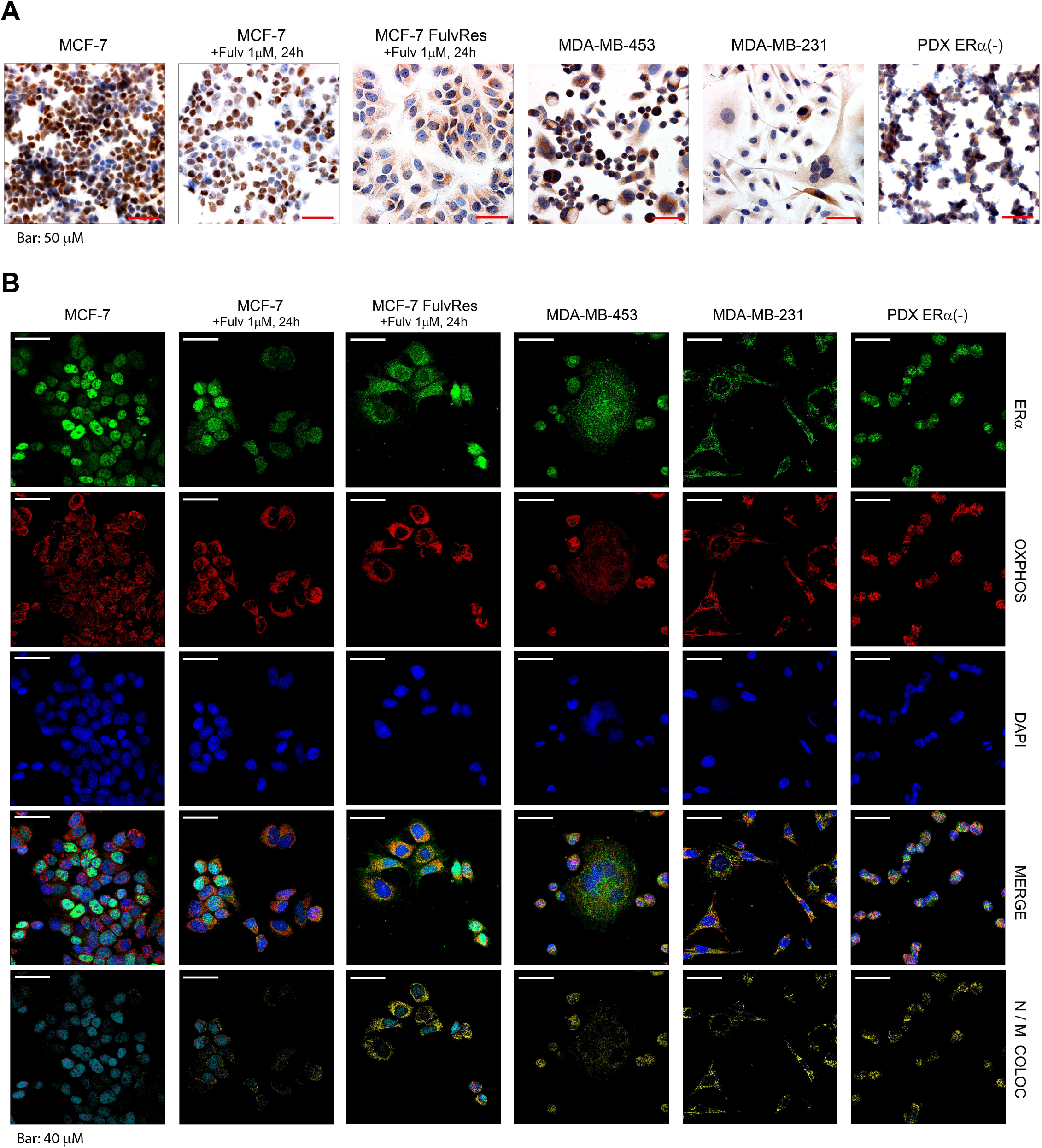
ERα-LBD localizes in the cytoplasm and mitochondria of breast cancer cells. **A,** Representative images of ERα staining by immunohistochemistry (IHC) in six different BC cell lines. All histological sections were counterstained with hematoxylin. Scale bar representing 50 μm is shown. Magnification: 40X. **B,** Representative confocal images of BC cells stained for ERα (green), mitochondria/OXPHOS (red) and nuclei/DAPI (blue). For each cell line, merged images and colocalization images are also shown. Color code for colocalization: cyan (ERα-Nuclei, N); yellow (ERα-Mitochondria, M). A scale bar for each image representing 40 μm is shown. Magnification: 63X.

Given the relative absence of ERα-LBD nuclear localization, we examined its role in canonical ERα signaling and we observed none of the transcriptional activation functions of full-length ERα. For instance, overexpression of ERα-LBD had no effect on Estrogen Response Element (ERE) luciferase expression or ERα target gene transcription in both ERα-FL(+) and TNBC cells (**Supplementary Fig. S2D** and **Fig. 3B-G**) (**26**).

To further establish the functional significance of cytoplasmic and mitochondrial ERα-LBD, we performed co-IP and mass spectrometry experiments aimed at identifying ERα-LBD protein-protein interactions (PPIs). We designed the experiment comparing MCF-7 cells overexpressing ERα-LBD with parental (NC), in the absence or presence of fulvestrant and we identified a total of 52 peptides associated with ERα-LBD (**Fig. 4A**; **Supplementary Fig. S2D**). Pathway enrichment and neural networking analyses performed on this protein set identified processes involved in carbohydrate metabolism (glycolysis and gluconeogenesis), cell signaling (MYC, mTOR/mTORC1, PIK3C1/AKT, ERα), hypoxia and angiogenesis (HIF1α and VEGFR signaling) and mitochondrial metabolism (oxphos and respiration) (**Fig. 4B** and **C**; **Supplementary Fig. S4A** and **Supplementary File S2**). Comparable experiments in TNBC models overexpressing or knocking down ERα-LBD identified a total of 93 peptides involved in ERα-LBD PPIs, and 29 of them were shared with those found in MCF-7 cells (**Fig. 4D**; **Supplementary Fig. S2D**). Similarly to the MCF-7 model, enriched pathways included signaling, carbohydrate and mitochondrial metabolism (including also fatty acid β-oxidation) and angiogenesis (**Fig. 4E** and **F**; **Supplementary Fig. S4B** and **Supplementary File S2**).

**Figure 4.**
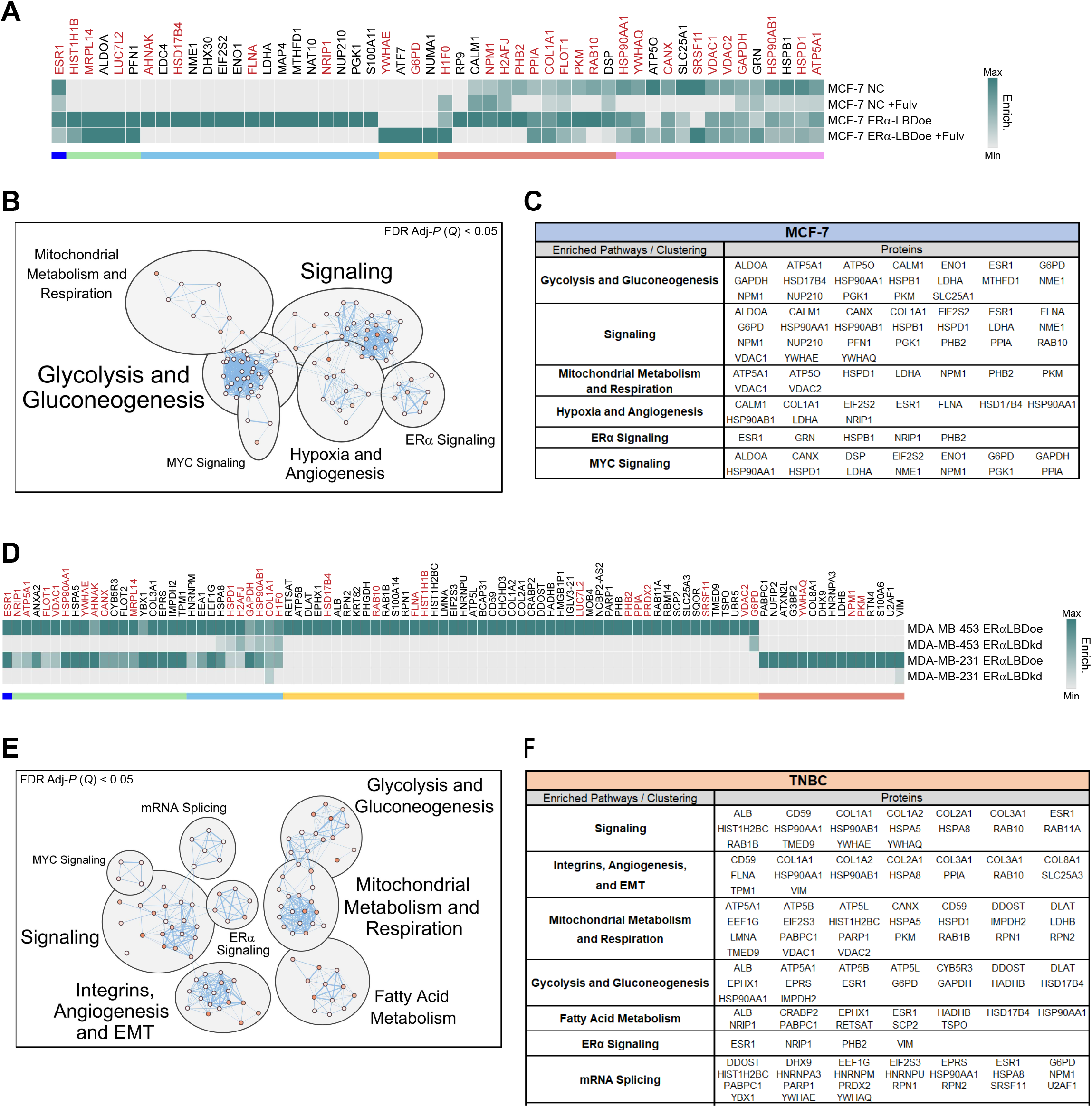
Predicted ERα-LBD network: protein-proteininteractions (PPls) and biological pathways. **A,** Heat map showing the enrichment level of proteins that preferentially bind ERα-LBD, based on ERα co-immunoprecipitation (Co­ IP) experiments followed by LC/MS analysis. Samples: NC and NC +Fulv (negative controls); ERα-LBDoe, ERα-LBDoe +Fulv (overexpression). Fulv = 1 μM fulvestrant treatment, 24 h. Different comparative criteria between samples were used to identify groups of ERα-LBD protein interactors, color-coded as follow: ERα (blue); proteins present only in ERα-LBD samples (green, cyan and yellow); proteins enriched more in ERα-LBD samples compared to NC, in the absence or presence of fulvestrant treatment (red and purple, respectively). Protein names labelled in dark red color are shared between MCF-7 and TNBC models (see below). **B,**’Enrichr’ platform based on multiple libraries was used to discover cell pathways in which ERα-LBD PPls are significantly involved (FDR *Adj-P* (*Q*) < 0.05), then clustered and annotated by using Cytoscape software and MCL algorithm. **C,** ERα-LBD PPls contribution to clustered pathways. **D,** Same as in **(A).** Samples: MDA-MB-453/-231 ERα-LBDoe (overexpression); MDA-MB-453/-231 ERα-LBDkd (knockdown). ERα PPls and protein groups, color-coded as follow: ERα (blue); proteins enriched in both TNBC cell lines overexpressing ERα-LBD, compared to ERα-LBD silenced cells (green and cyan); enriched in MDA-MB-453 ERα-LBDoe cells only (yellow); enriched in MDA-MB-231 ERα-LBDoe cells only (red). **E** and **F,** Same as in **(B** and **C)**.

### ERα-LBD regulates cell metabolism

The localization and protein-protein interaction analyses point to a potential metabolic function for ERα-LBD. We speculated ERα-LBD may play a role regulating glycolysis and mitochondrial respiration and so assessed the effect of ERα-LBD overexpression or knockdown on key metabolic parameters. In ERα-FL expressing MCF-7, control (NC) and ERα-LBDoe cells showed similar OCR (oxygen consumption rate, index for respiration) and ECAR (extracellular acidification rate, index for glycolysis) under basal conditions. However, following fulvestrant-mediated depletion of ERα-FL and stabilization of ERα-LBD, we observed significantly higher respiratory parameters (ATP production, basal and maximal respiration and spare respiratory capacity) and higher glycolytic parameters (glycolytic capacity and glycolytic reserve) (**Fig. 5A**; **Supplementary Fig. S5A**). In the TNBC models featuring knockdown of ERα-LBD, ERα-LBDkd cells were characterized by reduced levels of respiratory parameters (ATP production, basal and maximal respiration), compared to controls (NC). Notably, ERα-LBDkd led to the impairment of glycolysis only in MDA-MB-231 (**Fig. 5B** and **C**; **Supplementary Fig. S5B** and **C**).

**Figure 5.**
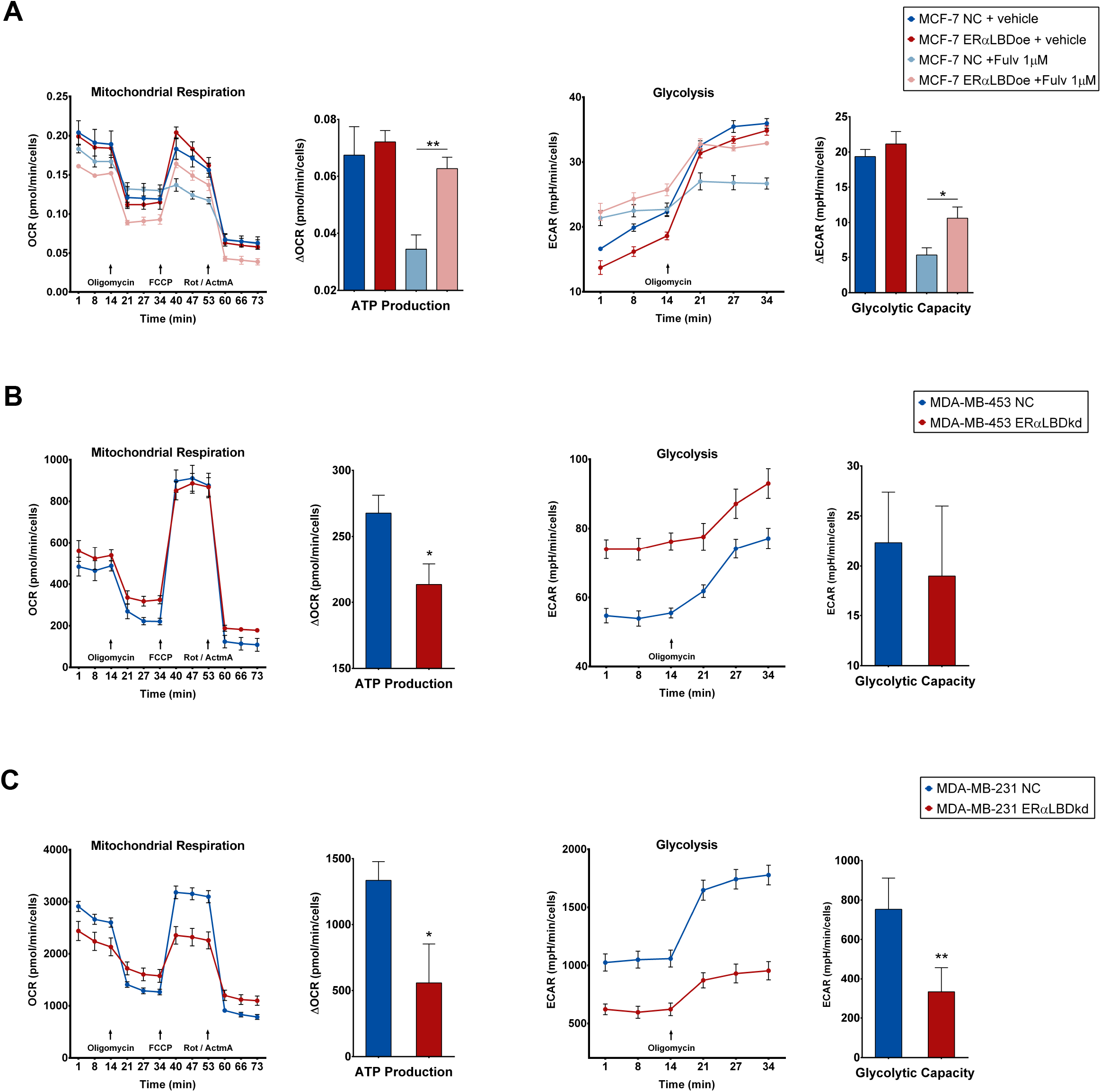
Effect of ERα-LBD ‘gain/ loss-of-function’ on BC cell metabolism. **A,** Mitochondrial respiration and glycolysis levels in MCF-7 cell clones. Fulvestrant 1 μM (24 h pre-treatment) or vehicle (DMSO) was added to cells. ERα-LBD overexpressing (ERα-LBDoe; in red) cells were compared to controls (NC; in blue). Analyses were carried out using XF Cell Mito Stress Kit (Agilent). Respiration was evaluated as oxygen consumption rate (OCR, pmol/min), normalized on cell number. Glycolysis was evaluated as extracellular acidification rate (ECAR, mpH/min), normalized on cell number. Calculation of metabolic parameters was based on t.OCR and t.ECAR values, following manufacturer’s guidelines. **B** and **C,** Mitochondrial respiration and glycolysis levels in MDA­ MB-453 and MDA-MB-231 cells. Cells with ERα-LBD knockdown (kd; in red) were compared to controls (NC; in blue). Analyses were carried out as described above. FCCP = p-trifluoromethoxy-phenylhydrazone; Rot= rotenone; AtmA = antimycinA. Data in the figures are presented as mean ± s.e.m. *(n* = 2 independent experiments). * *P* < 0.05, ** *P* < 0.01; unpaired t test, one-sided.

### ERα-LBD promotes growth and fulvestrant resistance, *in vitro* and *in vivo*

Given the impact of ERα-LBD on breast cancer cell metabolism, we investigated the effects of ERα-LBD on cell proliferation. *In vitro*, MCF-7 cells overexpressing ERα-LBD, compared to control cells (NC), showed no significant differences in growth. However, upon treatment with fulvestrant, control cells were growth inhibited while ERα-LBDoe cells continued to proliferate establishing a role for ERα-LBD in mediating fulvestrant resistance (**Fig. 6A**). Moreover, growth of MCF-7 FulvRes cells was potently impaired after ERα-LDB knockdown, especially under fulvestrant treatment (**Fig. 6B**). Beyond the effects in ERα-FL(+) cells, we also observed effects of ERα-LBD in TNBC cells. In both MDA-MB-453 and MDA-MB-231 we observed ERα-LBD knockdown to slow cell proliferation (~50%) (**Fig. 6C** and **D**).

**Figure 6.**
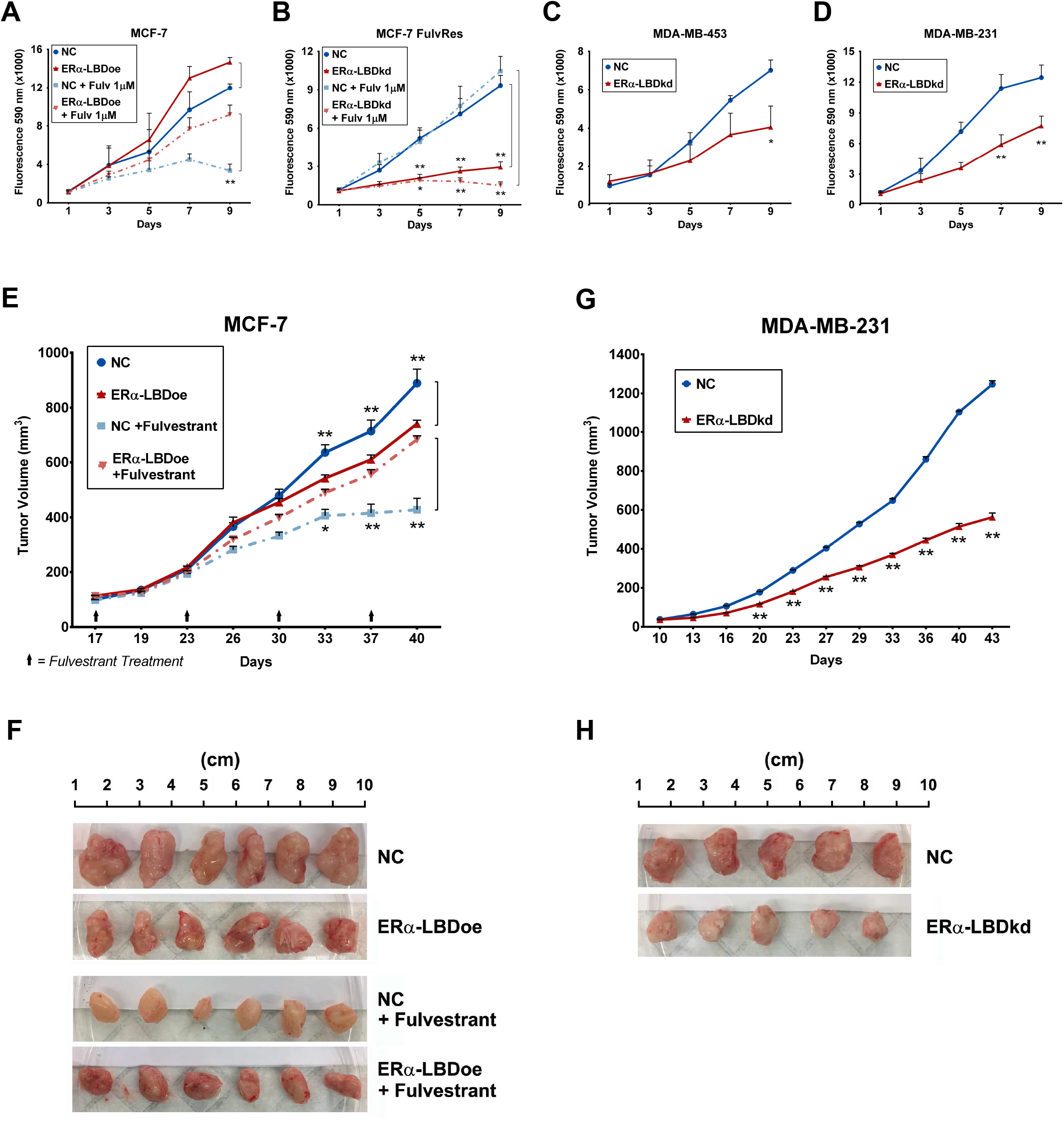
ERα-LBD promotes *in vitro* and *in vivo* growth and fulvestrant resistance. **A-D,** *In vitro* proliferation of BC cell clones, using resazurin reagent and expressed as fluorescence intensity (absorbance at 590 nm). ERα-LBD overexpression **(A)** or knockdown **(B-D)** was compared to controls (NC). In **(A)** and **(B),** cell proliferation was also tested in the presence or absence of fulvestrant 1 μM treatment *(n* = 3 independent experiments). **E-H,** Xenografts of BC cells implanted into the mammary fat pads of NOD scid gamma (NSG) mice. Plots in **(E)** and **(F)** show mean tumor volumes (mm^3^) as a function of time (days). MCF-7 model: when tumors reached 100 mm3, mice were randomized *(n* = 6/group) to weekly treatments with 200 mg/kg fulvestrant injected intramuscularly, and compared to control groups *(n* = 6/group). MDA-MB-231 model: *n* = 5/group. **F-H,** Images and size of tumors, isolated from animals after reaching endpoint. In **(A)** and **(E),** NC is in blue and ERα-LBD overexpression in red. In **(B-D)** and **(G),** NC is in blue and ERα-LBD knockdown in red. All data in the figure are presented as mean± s.e.m., ns = not significant,* *P* < 0.05, ** *P* < 0.01, two-way ANOVA.

To further verify the biologic significance of these effects *in vivo*, these models were grown orthotopically as xenograft models. As observed *in vitro*, MCF-7 ERα-LBDoe tumors continued to grow upon fulvestrant treatment while tumors from NC cells were expectedly growth inhibited with fulvestrant (**Fig. 6E** and **F**). Moreover, *in vivo* tumor growth for MDA-MB-231 ERα-LBDkd mice group was markedly reduced, when compared to NC group (**Fig. 6G** and **H**).

MCF-7 ERα-LBDoe cells showed a growth advantage when cultured in low-attachment (3-D growth) either in the presence or absence of fulvestrant treatment (**Supplementary Fig. S6A**), whereas ERα-LBD knockdown led to reduced 3-D growth in both MDA-MB-453 and −231 cells (**Supplementary Fig. S6B**). Similarly, the migratory capacity of BC models was enhanced in ER LBD overexpression models and impaired in knockdown models (**Supplementary Fig. S6C** and **D**).

### ERα-LBD expression is associated with proliferation, endocrine resistance, stemness and metabolism of breast cancer cells

Given the growth advantage, drug resistance and metabolic phenotypes afforded by ERα-LBD, we further evaluated its impact by gene expression analysis. We considered fulvestrant treated MCF-7 ERα-LBDoe and MCF-7 FulvRes cells as ‘gain-of-function’ models compared to MCF-7 control cells (NC). RNA-seq analysis led to the identification of ~17K differentially regulated genes (**Fig. 7A**; **Supplementary File S3**). Gene set enrichment analysis (GSEA) was performed on these same samples using specific gene set collections related to breast cancer, cell proliferation and cell metabolism (**Supplementary File S4**). Interestingly, gene sets involving cell signaling, breast cancer malignancy/endocrine resistance, EMT and stemness, carbohydrate and mitochondrial metabolism were found to be enriched in the ‘gain-of-function’ models (**Fig. 7B** and **C**; **Supplementary File S3**). Conversely, RNA-seq analysis of MDA-MB-453/-231 ERα-LBDkd “loss-of-function” models identified ~16K differentially regulated genes in TNBC ERα-LBDkd cells compared to NC cells (**Fig. 7d**; **Supplementary File S3**). Similarly to the MCF-7 model, gene sets associated with cell proliferation and survival, cell stemness and metabolism were found to be enriched in TNBC NC cells compared to TNBC ERα-LBDkd (**Fig. 7E** and **F**; **Supplementary File S3**).

**Figure 7.**
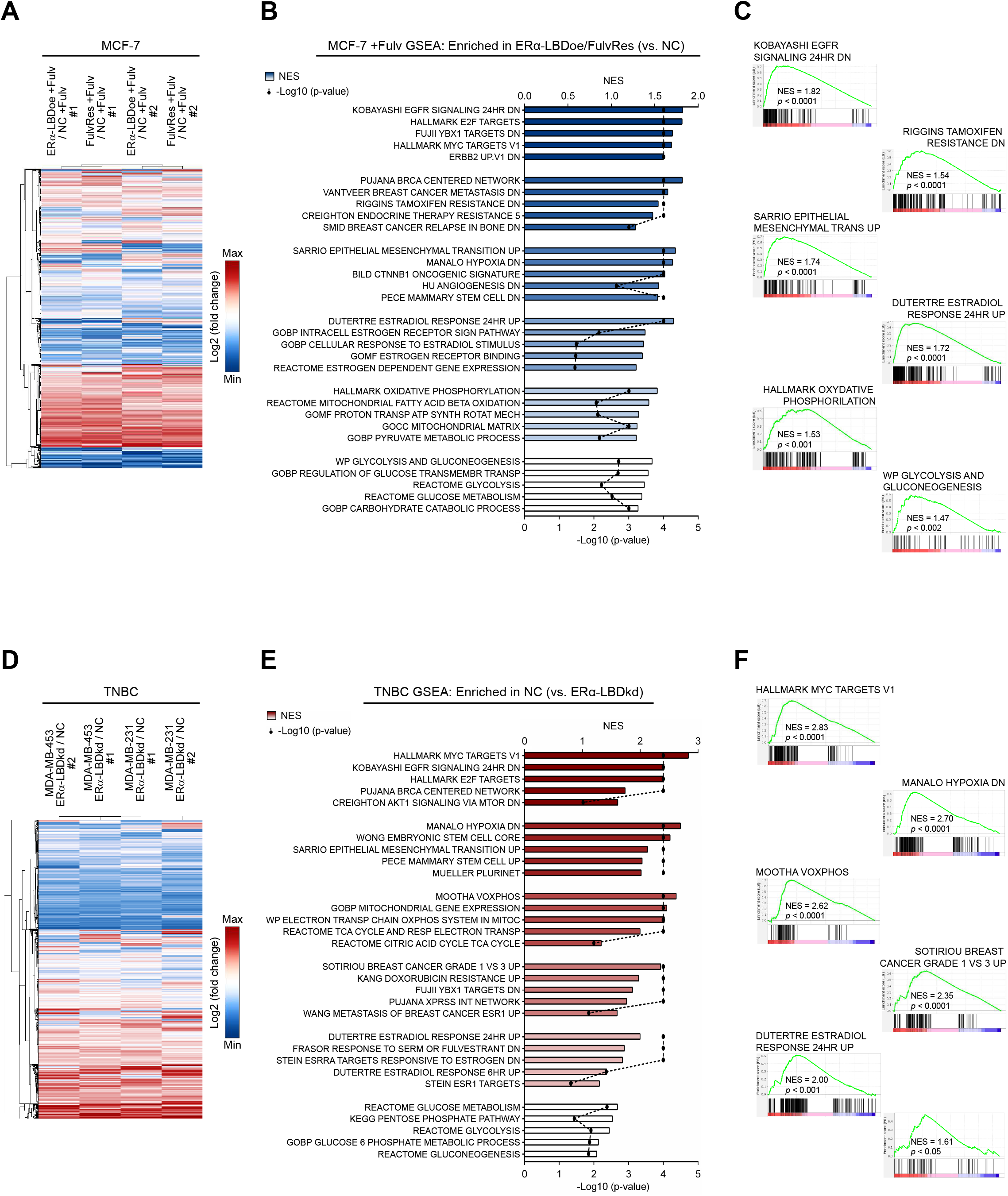
ERα-LBD expression is associated with proliferation, endocrine resistance, sternness and metabolism of breast cancer cells. **A-F,** Two different BC cell models were used to investigate cell phenotypes associated with ERα-LBD ‘gain/ loss-of-function’. **A-C** MCF-7 model, with 24 h fulvestrant 1 μM treatment (+Fulv): MCF-7 ERα-LBDoe, MCF-7 FulvRes and MCF-7 NC (control). **D-F** TNBC model: MDA-MB-453/-231 ERα-LBDkd (knockdown) and MDA-MB-453/-231 NC (controls). RNA samples were extracted from each cell line and analyzed by RNA-Seq *(n* = 3/sample). **A,** Heat map showing differentially expressed genes in MCF-7 ERα-LBDoe (+Fulv) and MCF-7 FulvRes (+Fulv), compared to MCF-7 NC. Gene expression (counts) was determined by using HTSeq and differential expression was calculated as log2(fold change). Only genes with >1 fold change were taken under consideration for the analysis. Hierarchical clustering based on Euclidean Distance was carried out. **B** GSEA analysis based on gene counts and showing gene sets that are both highly and significantly (*P* < 0.05) enriched in MCF-7 ERα-LBDoe (+Fulv) and MCF-7 FulvRes (+Fulv), compared to controls. NES = normalized enrichment score. Enriched gene sets were grouped and color-coded considering their association with specific gene set collections. Please refer to Methods section for details. **C,** Enrichment plots of interest, derived from GSEA analysis described in **(B). D,** Heat map showing differentially expressed genes in TNBC ERα-LBDkd cells, compared to controls (NC). Calculation and analysis was carried out as described in **(A). E,** GSEA analysis showing gene sets that are both highly and significantly (*P* < 0.05) enriched in TNBC NC samples compared to TNBC ERα-LBDkd, grouped/color-coded as in **(B). F,** Enrichment plots of interest, derived from GSEA analysis described in **(E).**

We further explored the role of ERα-LBD in breast cancer stemness and endocrine resistance. Cells from MCF-7 and MCF-7 FulvRes xenografts tumors were screened by flow cytometry and qPCR to measure the expression of CD44, SOX2 and SOX9, markers associated with stemness and endocrine resistance (**27**, **28**). MCF-7 FulvRes tumors were characterized by a higher percentage of CD44^High^ cells which showed higher expression of ESR1-LBD, SOX2 and SOX9, suggesting a correlation between ERα-LBD expression and stemness (**Supplementary Fig. S7B** and **C**).

### ESR1-LBD expression is identified in human breast cancers

Our data supports a role for ERα-LBD in fulvestrant resistance and proliferation of BC cells. To determine the clinical relevance of these observations, we examined ESR1-LBD expression by analyzing RNA-seq samples from human BC specimens.

ESR1-FL overall expression was assessed on the following collection of samples: ERα(+) BC primary tumors (*n* = 42); uninvolved breast tissue adjacent to ERα(+) primary tumors (*n* = 30); TNBC primary tumors (*n* = 42); uninvolved breast tissue adjacent to TNBC primary tumors (*n* = 21); metaplastic BC primary tumors (MpBC, *n* = 17) (**29**, **30**). As expected, ESR1-FL transcript was found to be markedly reduced in TNBC and MpBC samples (**Fig. 8A**). Notably, ERα expression is known to be present in normal breast epithelium (**31**). For each cancer sample, ESR1 exon expression was also calculated and, similarly to what we observed with BC cell lines (**Fig. 2A**), TNBC and MpBC samples showed higher amounts of 5’UTR exon 3a and CDS exons #4-8, compared to #1-3 (**Fig. 8B**). We also examined the levels of the ESR1-LBD transcript in BC groups, relative to ESR1-FL (**Fig. 8C**). 76% of ERα(+), 50% of TNBC and 81% of MpBC were characterized by high ESR1-LBD levels (Q3+Q4 quartiles, above the median calculated on all samples). Interestingly, only 17% of ERα(+)^ADJ^ and none of TNBC^ADJ^ samples were found to have high ESR1-LBD expression (**Fig. 8D**; **Supplementary Fig. S8A**).

**Figure 8.**
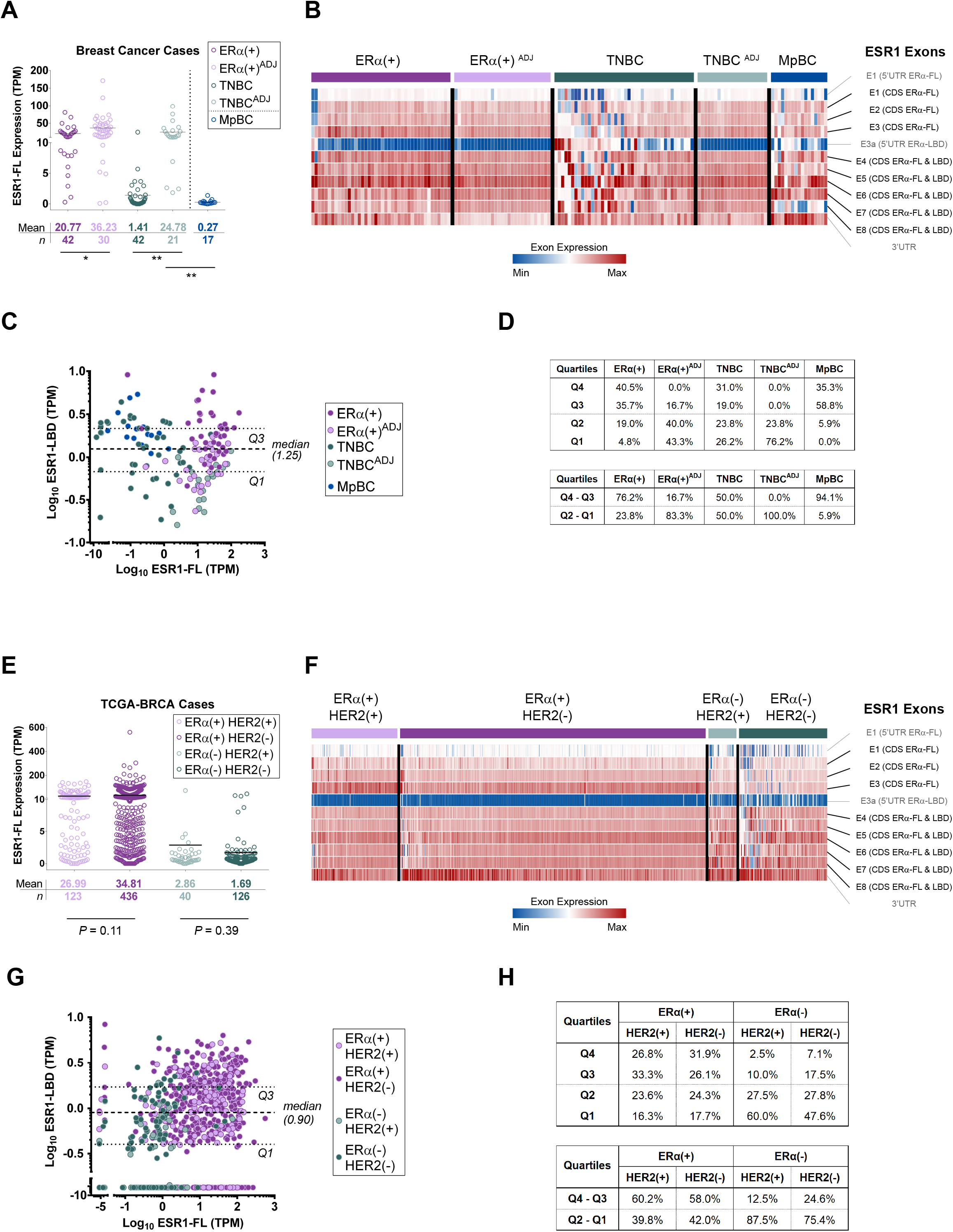
ESR1-LBD expression is identified in human breast cancers. **A,** RNA-seq samples from BC patients were grouped as follow: ERα(+) BC primary tumors *(n* = 42; in purple); uninvolved breast tissue adjacent to ERα(+) primary tumors (ERα+ADJ, *n* = 30; in light purple); TNBC primary tumors *(n* = 42; in teal); uninvolved breast tissue adjacent to TNBC primary tumors (TNBC^ADJ^, *n* = 21; in light teal); metaplastic BC primary tumors (MpBC, *n* = 17; in blue). For each group/sample, ESR1-FL transcript expression was calculated by using Kallisto and plotted as Transcripts Per Kilobase Million (TPM). **B,** Heat map showing ESR1 exon expression (including 5’/3’-UTR regions) in BC patient groups. Exon expression was calculated by using DEXSeq-count tool [# of read counts/exon length] and scaled by column. For each group, columns represent BC samples and were sorted by ESR1-FL expression (calculated as mean of all CDS exons), increasing from the left to the right side of the maps. CDS exons are indicated in black and untranslated regions (UTR) in grey. **C,** Dispersion plot for ESR1 transcript variants expression (ESR1-FL vs. ESR1-LBD), calculated as TPM. Dashed lines corresponding to ESR1-LBD expression median and Q1-Q3 quartiles are shown. Color code as described in **(A). D,** Percentage of samples from each group, distributed into quartiles. Percentage of samples above the median (Q4+Q3) and below the median (Q2+Q1) is also shown. **E,** RNA-seq samples from Breast Invasive Carcinoma study (TCGA-BRCA) were assigned to four different groups, based on ERα and HER2 IHC staining: ERα(+)/HER2(+) *(n* = 123; in light purple), ERα(+)/HER2(−) *(n* = 436; in purple), ERα(−)/HER2(+)*(n* = 40; in light teal) and ERα(−)/HER2(−) *(n* = 126; in teal). ESR1-FL transcript levels were calculated and plotted as described in **(A). F, G** and **H,** As described in **B, C** and **D**(respectively).* *P* < 0.05, ** *P* < 0.01, unpaired t test, two-sided.

Our second investigation was performed on TCGA-BRCA (The Cancer Genome Atlas Breast Invasive Carcinoma) RNA-seq samples (**32**). Based on the available IHC data, samples were grouped as: ERα(+)/HER2(+) (*n* = 123), ERα(+)/HER2(−) (*n* = 436), ERα(−)/HER2(+) (*n* = 40) and ERα(−)/HER2(−) (*n* = 126). The analysis of ESR1-FL expression showed low levels in ERα(−) samples but no significant differences between HER2(+) and HER2(−) cancers, in both ERα(+) and ERα(−) groups (**Fig. 8E**). As above, ESR1 exon expression is shown by using heat maps revealing a slightly higher abundance of exons #3a and #4-8, compared to #1-3 in the ERα(−) samples (**Fig. 8F**). ESR1-LBD transcript expression was also assessed in all groups and correlated to ESR1-FL (**Fig. 8G**). Higher ESR1-LBD levels (Q3+Q4) were identified in 60% of ERα(+)/HER2(+), 58% of ERα(+)/HER2(−), 13% of ERα(−)/HER2(+) and 25% of ERα(−)/HER2(−) (**Fig. 8H**; **Supplementary Fig. S8B**). All expression data are summarized in **Supplementary Files S5** and **S6**.

## DISCUSSION

Fulvestrant resistant breast cancer is frequently observed in the metastatic setting however little data exists on the molecular events underlying resistance (**13**, **20**, **33**, **34**). Our analysis of fulvestrant resistant BC models led to the identification of a novel ERα protein variant (ERα-LBD, 37.3 kDa MW) lacking the N-terminal transcriptional activation (AF1) and DNA binding domains (DBD) but including a portion of the hinge domain followed by the C-terminal domains LBD (Ligand Binding Domain) and AF2. Not only was ERα-LBD identified in fulvestrant resistant cell lines but also in TNBC models. Unlike full-length ERα which is degraded by fulvestrant, ERα-LBD is stabilized by this agent along with other ERα ligands such as estradiol and tamoxifen, leading to increased ERα-LBD expression (**Fig. 1**). We hypothesize that low expression of ERα-LBD in the absence of high levels of ligands may explain why this isoform has not been previously observed in BC cells by others. We also suggest that this feature should be exploited to better understand ERα protein stability and discover novel pharmacologic inhibitors of ERa and its relevant isoforms. A 37 kDa ER species expressed at low levels was previously isolated from the mouse uterus. Interestingly, this ER variant appeared to have the ligand-binding region and a portion of the hinge domain, but lacked a DNA-binding region (**35**).

The identity of ESR1-LBD transcript was determined by analyzing available sequence databases (specifically, FANTOM CAT) and by RNA-seq experiments, suggesting that the ESR1-LBD transcript might utilize a novel transcription start site and 5’UTR rather than processed by alternative splicing of the full-length transcript (ESR1-FL). The hypothesis of a new putative *ESR1* promoter was supported by preliminary data (not shown), describing potential transcriptional activity in the upstream genomic region from exon E3a. Additionally, we noted reduced expression of the 3’UTR sequence of ESR1-LBD in the ERα(−)/TNBC models, a phenomenon that could alter mechanisms of post-transcriptional inhibition mediated for example by microRNAs. Indeed, we demonstrated that ESR1-LBD is more stable than ESR1-FL and this may explain how a relatively low abundant transcript is translated into a detectable protein (**Fig. 2**). ESR1-LBD variant has not been previously described and we suggest that the shortening of the 3’UTR might account for this, as it might limit its detection in experiments based on poly(A)-enriched RNA samples.

The putative function of ERα-LBD was first investigated by examining its localization demonstrating its absence from the nucleus and enrichment in the cytoplasm and mitochondria of breast cancer cells (**Fig. 3**). We examined ERα-LBD protein-protein interactions by co-IP/MS using overexpression and knockdown models and determined a role in glycolysis/gluconeogenesis, mitochondrial metabolism, signaling and angiogenesis (**Fig. 4**). These observations were further supported by functional analyses revealing metabolic and cell/tumor growth advantages associated with ERα-LBD (**Figs. 5** and **6**) and RNA-seq analysis (**Fig. 7**).

An extra-nuclear role for ERα and its isoform ERα-36 has been well established most notably by their presence in the plasma membrane and their association with receptor tyrosine kinases and MAPK (**36**, **37**). Our data would suggest a distinct role for ERα-LBD as no apparent association with RTKs nor MAPK/ERK was identified. The role of mitochondrial ERα has also been described functioning as a transcription factor interacting with mtDNA and preventing UV-induced apoptosis as well as directly interacting with the mitochondrial protein HADHB (**4**, **38**). Although ERα-LBD has no DNA binding capacity, we also found HADHB among its protein partners, suggesting both unique and overlapping functions for ERα and ERα-LBD in the mitochondria (**Fig. 4**).

Additionally, a role of ERα-LBD in breast cancer stemness was suggested by RNA-seq analysis and qPCR demonstrating an enrichment of the ESR1-LBD transcript in fulvestrant resistant CD44^High^ tumor cells (**Fig. 7** and **Supplementary Fig. S7**). Importantly, ERα-LBD’s role in fulvestrant resistance was also demonstrated by overexpression studies on BC cell growth (**Fig. 6**) and supported by RNA-seq analysis (**Fig. 7**). Given the role of stemness with endocrine resistance, we suggest that ERα-LBD may provide a link between these phenomena (**13**, **14**, **39**, **40**). Furthermore, the higher abundance of this transcript and protein in cancer stem cells may also explain its relatively low detection in bulk tumor and cell line analyses.

Notably, some of ERα-LBD features are shared with the more recently discovered ERα isoform, ERα-36 (**24**, **25**, **41**). No cross-detection between ERα-LBD and ERα-36 was found in our models. Specifically, the C-terminal antibody we used for IF, IHC and co-IP studies does not recognize ERα-36. Additionally, the use of primers and an antibody specific for ERα-36 did not lead to its detection in our hands (**Supplementary Fig. S2**). We suggest that isoforms of ERα may play unique and/or overlapping critical roles in endocrine resistant disease, TNBC and prognosis. Identifying the unique contributions of these isoforms in breast cancer will be of interest.

Lastly, we demonstrated the presence of the ESR1-LBD transcript in primary BC specimens using publicly available RNA-Seq data (**29**, **30**, **32**). ESR1-LBD was absent or poorly expressed in non-malignant breast tissue but found to variable levels in ERα(+) tumors, TNBC and metaplastic breast cancers (MpBC). Remarkably, high levels of ESR1-LBD were identified in 16/17 of metaplastic breast cancers which is one of the most aggressive and chemo-resistant subtypes. The TCGA-BRCA study allowed us to examine the contribution of HER2 expression on ESR1-LBD expression. We observed higher levels of ESR1-LBD in TNBC (25%) as compared to ER-/HER2+ samples (13%), demonstrating differential expression of this transcript in the different subtypes of breast cancer (**Fig. 8**). Unfortunately, data from BC cases with acquired fulvestrant resistance were not available but we hypothesize that expression of ESR1/ERα-LBD would be found in a subset of these cancers. A limitation of our study is the lack of a selective ERα-LBD protein assay to test for this isoform in patient samples and thus cannot ascertain its protein expression and prevalence.

In summary, we have identified a novel ERα isoform “ERα-LBD” in ERα(+) fulvestrant resistant cancer cells and TNBCs. The development of specific strategies for ERα-LBD detection (*e.g.* antibodies, probes) is essential. Further studies are ongoing, to unravel a broader presence and the unique role that ERα-LBD plays in tumorigenesis and cancer progression, as well as its prognostic significance.

## METHODS

### Cell culture and reagents

All cell lines were obtained from ATCC (American Type Culture Collection) and authenticated using short tandem repeat (STR) analysis. Cells were cultured in MEM or RPMI-1640 medium (MSKCC Media Prep, USA) supplemented with 10% heat-inactivated fetal bovine serum (Gibco, USA) and 1% penicillin/streptomycin (Gibco, USA) and maintained at 37°C and 5% CO_2_ in humidified atmosphere. Reagents used for cell culture and treatments: fulvestrant (Selleckchem, USA), SNX-2112 (Selleckchem, USA), charcoal stripped fetal bovine serum (CSS, Gibco, USA), 17β-estradiol (E2, Sigma-Aldrich, USA), 4-Hydroxytamoxifen (4-OHT, Sigma-Aldrich, USA). Further information on cell lines and clones in **Supplementary Data**.

### Western blot

Cells were homogenized in RIPA lysis buffer for protein extraction, supplemented with protease inhibitors (Thermo Scientific, USA). Denatured proteins were separated in 12% SDS–PAGE and then transferred onto nitrocellulose papers (Pall, USA). After blotting, nitrocellulose papers were incubated with specific antibodies. Primary antibodies: anti-ERα F-10 (Santa Cruz Biotechnology, USA, 1:500); anti-β-actin 13E5 (Cell Signaling Technology, USA, 1:2000). Secondary antibodies (HRP conjugated): anti-mouse (Cytiva, USA, 1:5000). Immunolabelling was visualized using ECL procedure (PerkinElmer, USA).

### Protein structure analysis

Protein sequence and features were analyzed using Sequence Analysis software (Informagen, USA). Three-dimensional protein models were generated by Phyre^2^ web portal (www.sbg.bio.ic.ac.uk) and visualized by using RasMol software (http://www.openrasmol.org).

### RNA capture-sequencing and ESR1 transcripts analysis

Total RNA was extracted as described elsewhere (**42**) and samples (100 ng) were input in the RNA library construction using the KAPA RNA Hyper library prep kit (Roche, Switzerland). Customized adapters with unique molecular indexes (UMI) (Integrated DNA Technologies, USA) and Sample-specific dual-indexes primers (Integrated DNA Technologies, US) were added to each library. Each RNA library was pooled for hybridization capture with customized ESR1 Panel (Integrated DNA Technologies, USA) using a capture protocol modified from NimbleGen SeqCap Target Enrichment system (Roche, Switzerland). Libraries were then sequenced on an Illumina MiniSeq with paired-end reads (x150 cycles, 1.4millions reads/sample). Raw sequencing data output was processed for expression analysis using STAR Aligner (**43**), DEXSeq-count (**44**), Cluster 3.0 and Java TreeView (**45**, **46**), DEXSeq and plotDEXSeq (**44**), Kallisto (**47**) (**Supplementary Fig. S2H**).

### Immunohistochemistry and confocal microscopy

ERα immunostaining (IHC) was performed on Benchmark Ultra using the ultraView DAB Detection kit (Ventana, USA). Antigen retrieval was performed onboard with UltraCC1 buffer (pH 8.2–8.5) at 95°C for 52 min. Primary antibody (anti-ERα F-10, Santa Cruz Biotechnology, USA) 1:100 for 28 min at 37°C. Secondary antibody 1:100 for 1 h. Images were obtained using a Zeiss Axiovert Widefield Microscope and Zeiss ZEN software (Carl Zeiss, Germany). Immunofluorescence was performed using Leica Bond RX stainer and the Bond Polymer Refine Detection kit (Leica, Germany). Antibodies and detection: OXPHOS (2μg/ml, Invitrogen, USA); ERα D8H8 (1μg/ml, Cell Signaling Technology, USA); Alexa Fluor Tyramide signal amplification reagents (Life Technologies, USA); 4’,6-diamidino-2-phenylindole (DAPI, Sigma-Aldrich, USA). Slides were mounted in Mowiol 4-88 (Calbiochem, USA). Confocal imaging was performed on a Leica SP8 inverted microscope (Leica, Germany). Image processing and analysis (2-D and 3-D) was performed using Imaris software (Bitplane, CH).

### ERα Co-Immunoprecipitation and LC/MS

Protein were extracted using Co-IP buffer (NaCl 150 mM, EDTA 0.5 mM, NP-40 0.5% (v/v), Tris-HCl 10 mM pH 7.4, PMSF 1 mM). Immunoprecipitation (IP): anti-ERα antibody D8H8 1:75 (Cell Signaling Technology, USA) and Dynabeads™ Protein A (Invitrogen, USA). IP samples were washed 4 times with 50 mM ammonium bicarbonate buffer and collected by centrifugation. Samples were then digested overnight with 2 μg trypsin in 80 μl of 50 mM ammonium bicarbonate at 37°C. Desalted and dried peptides were reconstituted in 10 μl 0.1% (vol/vol) formic acid and analyzed (4 μl) by microcapillary liquid chromatography with tandem mass spectrometry. Further details in **Supplementary Methods**.

Pathway enrichment analysis was performed by using Enrichr platform (https://maayanlab.cloud/Enrichr). Enriched pathways network, clustering and protein– protein interaction (PPI) network analyses were carried out using Cytoscape, EnrichmentMap, AutoAnnotate and stringApp software (**48**).

### Metabolic assay

Mitochondrial respiration and glycolysis were assessed using Seahorse extracellular flux analyzer (XFe96) and Seahorse XF Cell Mito Stress Test Kit (Agilent Technologies, USA). Cells were seeded in 96-well plates (2 × 10^4^ cells/well), with or without Fulvestrant 1 μM depending on the experiments. Raw data output was collected and analyzed using Wave Software (Agilent Technologies). Procedures and OCR/ECAR data interpretation were carried out accordingly to manufacturer’s guidance.

### Animal models

NOD scid gamma (NSG) female mice at age of 6-8 weeks were obtained from Jackson Laboratory (USA) and maintained in pressurized ventilated caging. To sustain tumor growth in MCF-7 models, 17β-estradiol pellets (0.18 mg) were implanted subcutaneously 3 days before BC cells injection. For both MCF-7 and MDA-MB-231 models, cancer cells were injected in the mammary fat pads (MFPs) of mice. For each experimental sample, cell suspensions were mixed with an equal volume of Matrigel (BD Biosciences, USA). For the MCF-7 model, injectable fulvestrant (Faslodex®, AstraZeneca, UK) was given intramuscularly in the tibialis posterior/popliteal muscles (100 mg/injection/once a week) for 20 days. Control mice received isotype control (placebo) or PBS injection. Tumor volumes were measured with vernier calipers starting from 14 days after cell implantation. Animals were sacrificed as they reached experimental endpoint. All procedures and experiments were completed in accordance with the Guidelines for the Care and Use of Laboratory Animals and were approved by the Institutional Animal Care and Use Committees at MSKCC (MSKCC#12-10-016).

### RNA sequencing

Total RNA was extracted from cell lines as described elsewhere (**42**) and sequenced on an Illumina HiSeq instrument (2×150 paired-end, 100 million reads/sample) by GENEWIZ, LLC. (USA). Raw sequencing data output was processed for downstream analyses using the following bioinformatic tools: STAR (reads alignment) (**43**), HTSeq (gene expression) (**49**), Cluster 3.0 and Java TreeView (hierarchical clustering and heat map) (**45**, **46**), GSEA (https://www.gsea-msigdb.org) (**Supplementary Fig. S7A**; **Supplementary File S4**).

### Breast cancer patients data

RNA-seq FASTQ files of 135 breast cancer were obtained from the European Nucleotide Archive, ENA (Study Accession: PRJNA251383; **Supplementary File S5**) (**29**). RNA-seq FASTQ files of 17 metaplastic breast cancer were kindly provided by Dr. Jorge S Reis-Filho (**Supplementary File S5**) (**30**). RNA-seq bam files of 1097 breast IDC patients were obtained from TCGA breast cancer (BRCA) cohort (TCGA dbGaP Accession ID: phs000178.v11.p8; **Supplementary File S6**) (**32**). Raw data were processed by using STAR (reads alignment) (**43**), DEXSeq-count (**44**), Kallisto (**47**) and Morpheus (https://software.broadinstitute.org/morpheus).

### Statistics

Data were expressed as mean ± s.e.m. Student’s t test (unpaired/paired; one/two-tailed) or Analysis of Variance (two-way ANOVA, followed by Fisher’s test or Sidak’s/Tukey’s correction for multiple comparisons) were used to assess the statistical significance of the differences. For DEXSeq analysis, *P* values were adjusted for multiple testing using Benjamini & Hochberg (BH) correction (FDR). For Enrichr analysis, *P* values were computed from the Fisher exact test and FDR adjusted *P* values (*Q* values) were used to filter enriched pathways for Cytoscape and Enrichment Map analysis. For statistics on GSEA analysis, please refer to official website (www.gsea-msigdb.org). Differences were considered statistically significant at *P* < 0.01 and *P* < 0.05.

## ACKNOWLEDGMENTS AND FUNDING SOURCES

This work was supported by grants from National Institute of Health to MSKCC’s Cancer Center (P30CA008748) and to S.C. (R01CA204999), and a grant from University of Bologna to E.S. (RFO 2015). J.B. acknowledges Charles and Marjorie Holloway Foundation, Sussman Family Fund, Lerner Foundation and Beth C. Tortolani Foundation. S.C. also acknowledges support by the Breast Cancer Research Foundation. F.P. is partially funded by a National Institutes of Health/ National Cancer Institute K12CA184746 grant.

