## Supplementary data, figures and methods for "ERα-LBD, a novel isoform of estrogen receptor alpha, promotes breast cancer proliferation and endocrine resistance"

#### SUPPLEMENTARY METHODS

##### Cell proliferation and manipulation

Cells (1000/well) were seeded into 96-well plates and, accordingly to experimental design, vehicle (DMSO 0.01%) or fulvestrant 1 µM was added to the medium. Media and drug were replaced every 3 days. At different time points (day 1, 3, 5, 7 and 9), 20 µl of Resazurin (R&D Systems, USA) was added to 180 µl of medium in each well and incubated for 4 h. Fluorescence (adsorbance 590 nm) was measured for each well using a SpectraMax M5 microplate reader (Molecular Devices, USA) and correlated to cell growth.

To generate fulvestrant-resistant MCF-7 (MCF-7 FR) cells, cells were cultured in the presence of increasing concentrations of fulvestrant (0.1 µM to 1 µM). Cells were deemed resistant when they grew as parental cells in 1 µM fulvestrant. MCF-7 Y537S CRISPR knock-in cells were generated as described elsewhere (**1**). PDX models of ERα(+)/(-) metastatic BC were generated and maintained as described elsewhere (**2**).

BC cell clones overexpressing ERα-LBD (ERα-LBDoe) were generated using stable retroviral transduction. Briefly, ERα-LBD coding sequence was cloned into pBABE-Puro vector (Addgene, USA; **Supplementary File S7**). To generate retroviral particles, three plasmids were co-transfected into 293T packaging cells using Lipofectamine 2000 (Invitrogen, USA): pBABE-Puro-ERαLBDoe, pCMV-VSV-G and pUMVC (Addgene, USA). Viral supernatants were collected 48 h and 72 h later, centrifuged to remove cell debris, filtered through 0.45-µm filters (Millipore, USA) then used with polybrene 8 µg/ml polybrene (Santa Cruz Biotechnology, USA) to transduce BC cell lines. After 2 days, stable clones were selected with puromycin 2 µg/ml.

BC cell clones with ERα-LBD knockdown (ERα-LBDkd) were generated using stable lentiviral transduction and CRISPR/CAS9 technology for genome editing. Briefly, three different guide RNA sequences targeting ESR1 gene (exon #6) were cloned in the BsmBI site of lentiCRISPRv2 vector (Addgene, USA; **Supplementary File S7**) to create a pool of pCR-ERαLBDkd lentivectors. To generate lentiviral particles, the following plasmids were co-transfected into 293T packaging cells: pCR-ERαLBDkd (pool), pCMV-VSV-G and psPAX2 (Addgene, USA). Infection and selection procedures: same as retroviral approach (see above).

##### Gel extraction and mass spectrometry

Protein lysates from MCF-7 FR and MDA-MB-231 (both treated with fulvestrant 1 µM for 24h) were obtained using IP buffer (NaCl 150 mM, EDTA 0.5 mM, NP-40 0.5% (v/v), Tris-HCl 10 mM pH 7.4, PMSF 1 mM) and immunoprecipitated using anti-ERα antibody (Cell Signaling Technology, USA). IP samples were run on 2 electrophoresis gels, one used for staining with SimplyBlue Safe Stain (Invitrogen, USA) and band extraction, the other for WB sample check. Please refer to 'Methods' section of the manuscript for IP and WB procedures. In-gel digestion was performed using the method by Shevchenko et al. (2006) (**3**). Briefly, gel bands were excised, washed with acetonitrile and 100 mM ammonium bicarbonate solution (1:1) for 30 min, dehydrated with 100% acetonitrile for 10 min until gel slices shrunk and excess acetonitrile was removed, then slices were dried in a speed-vac for 10 min without heat. Gel slices were reduced with 5 mM DTT for 30 min at 56°C in a thermo-mixer with gentle mixing, removed, allowed to cool to room temperature then alkylated with 11 mM IAA for 30 min in the dark. Gel slices were washed with 100 mM ammonium bicarbonate and 100% acetonitrile for 10 min each. Excess acetonitrile was removed and the slices dried in a speed-vac for 10 min without heating. Gel slices were then rehydrated in a solution of 25 ng/ml trypsin in 50 mM ammonium bicarbonate on ice for 30 min. Digestions were performed overnight at 37°C in a thermo-mixer with gentle mixing. Digested peptides were collected and further extracted from gel slices in extraction buffer (5% formic acid and 50% acetonitrile, 1:2 vol/vol) at high speed mixing. Extractions were combined and dried down in a vacuum centrifuge. Peptides were desalted with C18 resin-packed stage-tips, lyophilized to dryness, then re-constituted in 3% acetonitrile/0.1% formic acid for LC-MS/MS analysis. LC-MS/MS Analysis LC-MS/MS was performed using a Waters NanoAcquity LC system (with a 100 mm inner diameter x 10 cm length C18 column (1.7 mm BEH130; Waters, USA) configured with a 180 mm x 2 cm trap column coupled to a Thermo Q-Exactive Plus orbitrap mass spectrometer (Thermo Scientific, USA). Trapping was performed at 15 ml/min 0.1% formic acid (Buffer A) for 1 min. The LC gradient was 0.5% to 50% B (100% acetonitrile; 0.1% formic acid) over 90 min at 300 nl/min. MS data were collected in data dependent acquisition (DDA) mode utilizing a top ten precursor ion selection for HCD fragmentation. Full MS scans were performed with the following parameters: Resolution: 70,000; AGC target: 1e6; Maximum IT: 50 ms; Scan Range: 400 to 1600 m/z. DDA parameters were as follows: Resolution: 17,500; AGC target 5e4; Maximum IT: 50 ms; Isolation window: 1.5 m/z; NCE: 27; Minimum AGC target: 2e3; Intensity Threshold: 4e4; Dynamic Exclusion: 15 s; Charge exclusion: unassigned, 1, 6-8, >8. All MS/MS samples were analyzed using Mascot (Matrix Science, UK). Mascot was set up to search the SwissProt_sprot_20170705_20180523 database (selected for Homo sapiens, unknown version, 20215 entries) assuming the digestion enzyme trypsin. Mascot was searched with a fragment ion mass tolerance of 0.080 Da and a parent ion tolerance of 10.0 PPM. Carbamidomethyl of cysteine was specified in Mascot as a fixed modification. Deamidated of asparagine and glutamine, oxidation of methionine, acetyl of the N-terminus and phosphorylation of serine, threonine and tyrosine were specified in Mascot as variable modifications. Mass spectrometry data were further processed using Scaffold software (Proteome Software Inc., USA). Peptide identifications were accepted if they could be established at greater than 90.0% probability to achieve an FDR less than 1.0% by the Peptide Prophet algorithm with Scaffold delta-mass correction. Protein identifications were accepted if they could be established at greater than 89.0% probability to achieve an FDR less than 1.0% and contained at least 1 identified peptide. Protein probabilities were assigned by the Protein Prophet algorithm. Proteins that contained similar peptides and could not be differentiated based on MS/MS analysis alone were grouped to satisfy the principles of parsimony. Proteins sharing significant peptide evidence were grouped into clusters. Protein sequence and features were analyzed using Sequence Analysis software (Informagen, USA). Three-dimensional protein models were generated by Phyre$^2$ web portal (www.sbg.bio.ic.ac.uk) (**4**).

#### Western blot

For the procedure, please refer to 'Methods' section of the manuscript. Primary antibodies used for supplemental experiments: anti-ERα 1D5 (Invitrogen, USA), anti-ERα-36 (Alpha

Diagnostic International, USA); anti-ERα D8H8 (Cell Signaling Technology, USA); anti-β-actin 13E5 (Cell Signaling Technology, USA). Secondary antibodies (HRP conjugated): anti-rabbit (Cell Signaling Technology, USA); anti-mouse (Cytiva, USA). Immunolabelling was visualized using ECL procedure (PerkinElmer, USA).

##### ESR1-LBD transcript variant prediction

Comprehensive ESR1 transcript variants annotation was obtained from Ensembl Genome Browser (www.ensembl.org). We used ZEMBU Genome Browser (https://fantom.gsc.riken.jp/zenbu/) for variant prediction study, showing data collected on ESR1 gene, based on FANTOM5 and FANTOM CAT analyses (**5**, **6**). For ESR1 gene and transcripts details please refer to this link:

https://fantom.gsc.riken.jp/zenbu/gLyphs/#config=eljPd0OTPhloifL81WStGC;loc=hg19::chr6:152118533..152432431+.

##### RNA extraction and real-time qPCR

Total RNA was extracted from cells using Trizol reagent (Invitrogen, USA), according to the manufacturer's instructions. Extracted RNA samples were quantified and then treated with DNase I to remove any genomic DNA contamination, using Ambion DNase I kit (Invitrogen, USA). Reverse transcription was carried out using iScript™ Select cDNA Synthesis Kit (Bio-Rad, USA). cDNA levels were analyzed by real-time PCR using TaqMan Universal PCR Master Mix or SYBR Select Master Mix reagents and ViiA 7 Real-Time PCR system, according to the manufacturer's instructions (Applied Biosystems, USA). Melting curve data were collected to check PCR specificity. Samples were run in triplicate and mRNA levels were normalized against those of RPLP0 or β-actin. Relative expressions were calculated using the formula $2^{-2\Delta Ct}$ values. RT-PCR was carried out by using OneTaq Hot Start 2X Master Mix (New England Biolabs, USA). PCR samples were separated on a 1.5% agarose gel and results visualized using Gel Doc XR+ Imaging System (Bio-Rad, USA). For RNA Decay Assay, BC cells were seeded in 6-well plates and treated with actinomycin-D 5 μg/ml (Sigma-Aldrich, USA). RNA samples were collected at different time points (0, 4, 8, 24, 48 h). All PCR primers are summarized in **Supplementary File S7**.

##### Transfection and luciferase assay

Cells were seeded in 24-well plate (2 x $10^5$ cells/well) 1 day before transfection and treatments were added accordingly to the experimental design (vehicle, DMSO 0.01% or fulvestrant 1 μM). Cells co-transfected with 3x-ERE-TATA-Luciferase reporter plasmid (750 ng) and pRL-TK Renilla Luciferase plasmid (75 ng) (Addgene, USA) by using Lipofectamine 2000 (Invitrogen, USA) and following reagent protocol. Cells were harvested 24 h after transfection and cell lysates were used for Dual-Luciferase® Reporter Assay System analysis, according to the manufacturer's instructions (Promega, USA). Luciferase bioluminescence measurements were performed with the Veritas™ Microplate Luminometer (Promega, USA). For each sample, Firefly luciferase activity was normalized against Renilla luciferase activity.

**Protein-protein interaction (PPI) network**

For procedures, please refer to the Methods section of the manuscript.

**Microscopy, 3-D growth and wound healing assay**

MCF-7 cells (WT and FulvRes) were seeded into 6-well plate (1 x $10^5$ cells/well) and treated accordingly to the experimental design. After controls reached 100% confluence, images of cells were captured with Zeiss Axiovert Microscope and processed with Zeiss ZEN software (Carl Zeiss, Germany). For 3-D growth, BC cell clones were suspended sparsely and plated in 24-well ultra-low attachment plates (Corning, USA), 3 x $10^3$ cells/well. Treatments (DMSO 0.01% or fulvestrant 1 µM) were added to medium accordingly to experimental design. After 5 days, representative images of cells were acquired with a Zeiss Axiovert Microscope. Quantification of cell 3-D proliferation was based on the optical density (OD 600 nm) of cell suspensions and determined using a SpectraMax M5 microplate reader (Molecular Devices, USA). For the wound healing assay, about 1x$10^5$ cells were seeded into each well of 12-well plates and cultured until 100% confluence. The 'scratch' was created with a p200 pipet tip on the cell monolayer through the center of the well. The debris was removed by washing the well with 1 ml of culture medium and then 1 ml of medium (with 1% serum) was added into each well. Treatments (DMSO 0.01% or fulvestrant 1 µM) were also added accordingly to experimental design. The plate was incubated at 37°C, and images of the 'scratches' were captured at various time points with a Zeiss Axiovert Microscope. The width of scratch was measured with FIJI software.

**Flow cytometry (FACS)**

For FACS/flow analyses, tumors were digested in sterile Epicult media (StemCell Technology, Canada), minced with sterile razor blades and incubated for 3 hours in the presence of collagenase/hyaluronidase (1,000 Units/sample). Cells were washed with sterile filtered PBS supplemented with 1% BSA (PBS-BSA 1%) and filtered through a 40 mm nylon mesh (BD Biosciences, USA). Cells were then stained in a volume of 100 µL (PBS-BSA 1%) with CD44-APC antibody (100 ng/$10^6$ –$10^8$ cells (IM7, eBiosciences, USA) on ice for 30 min and analyzed by flow cytometry at the MSKCC's flow core with BD FACS Aria I instrument (BD Biosciences, USA). Samples were analyzed for cell population distribution and sorted for viability (DAPI$^{neg}$) and CD44 expression. For flow plot analyses, samples were run using FlowJo 7.5 software (Tree Star, USA).

**Breast cancer patients data**

For procedures, please refer to the Methods section of the manuscript.

**Statistics**

For analyses, please refer to the Methods section of the manuscript.

### Supplementary Fig. S1

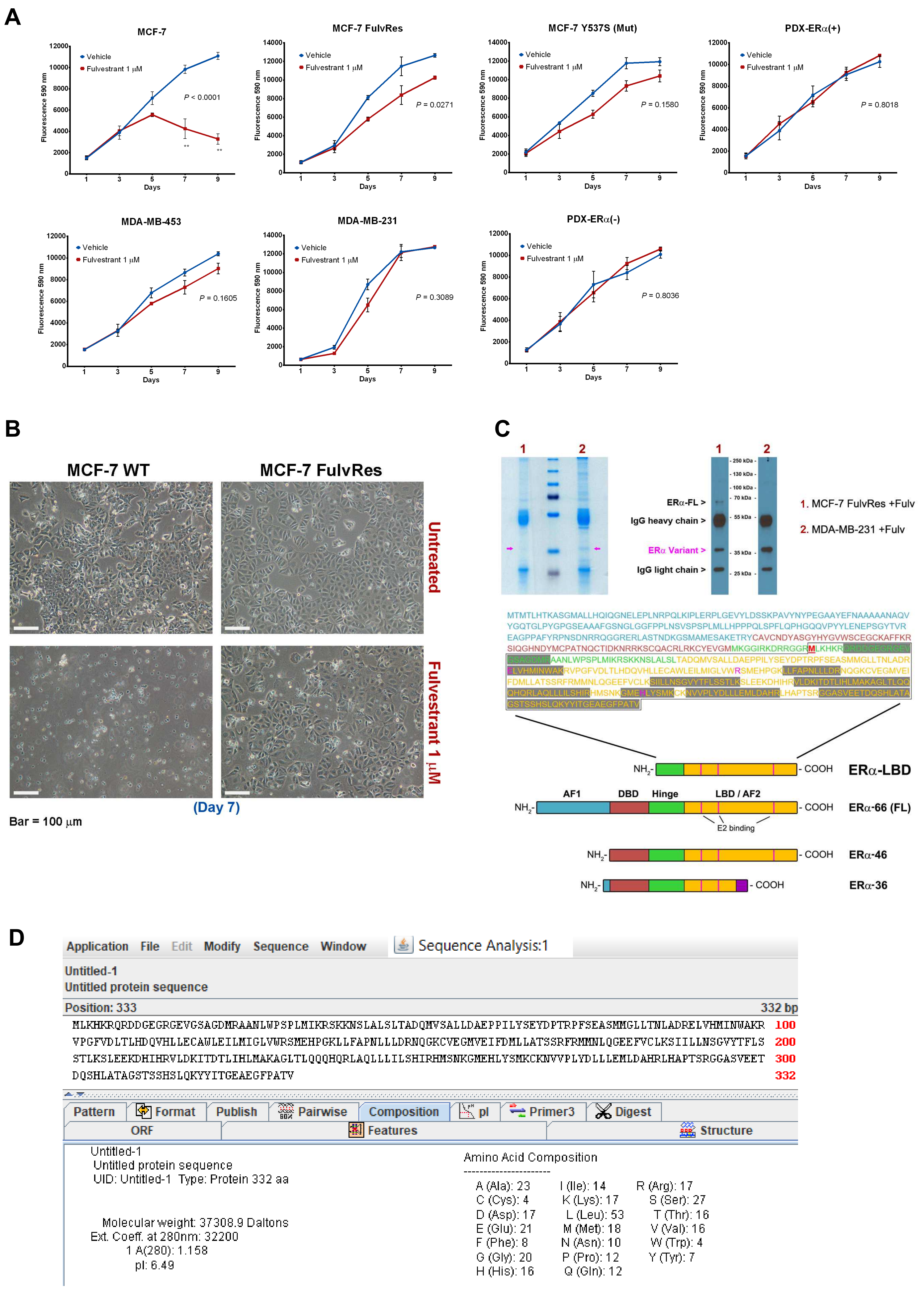

E

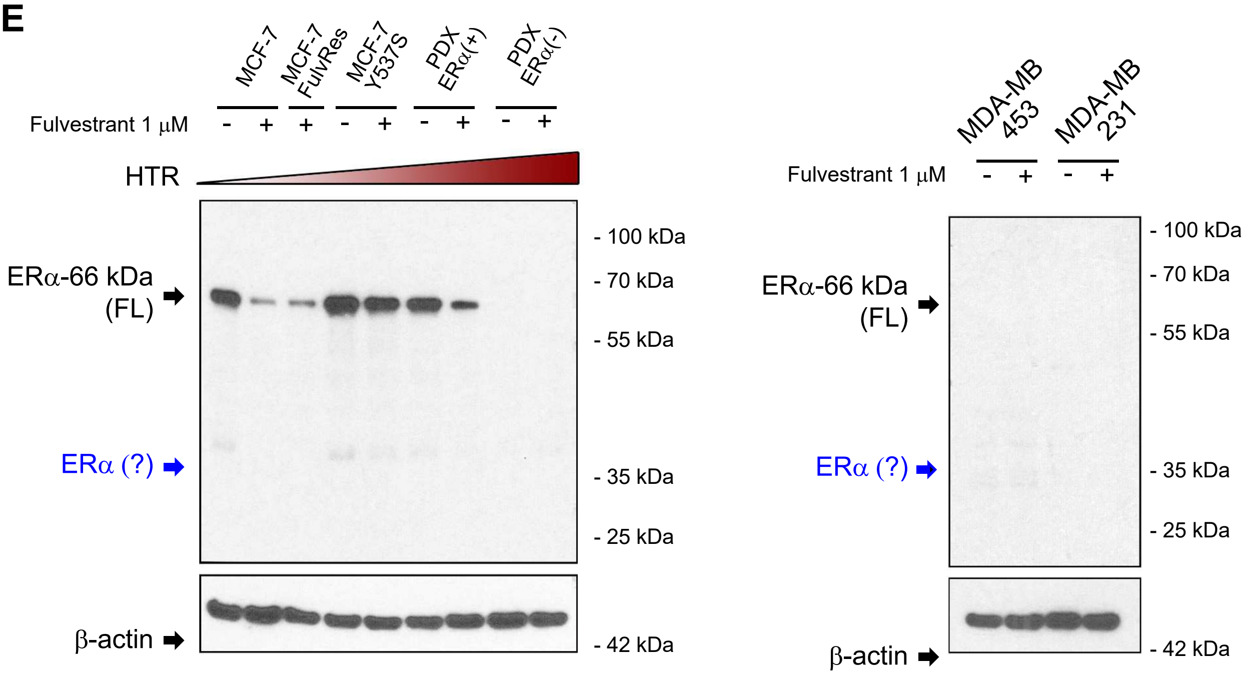

MCF-7
MCF-7 FulvRes
MCF-7 Y537S
PDX ERα(+)
PDX ERα(-)
Fulvestrant 1 µM
- + + - + - + - +
HTR
ERα-66 kDa (FL)
ERα (?)
β-actin
- 100 kDa
- 70 kDa
- 55 kDa
- 35 kDa
- 25 kDa
- 42 kDa
MDA-MB 453
MDA-MB 231
Fulvestrant 1 µM
- + - +
ERα-66 kDa (FL)
ERα (?)
β-actin
- 100 kDa
- 70 kDa
- 55 kDa
- 35 kDa
- 25 kDa
- 42 kDa


F

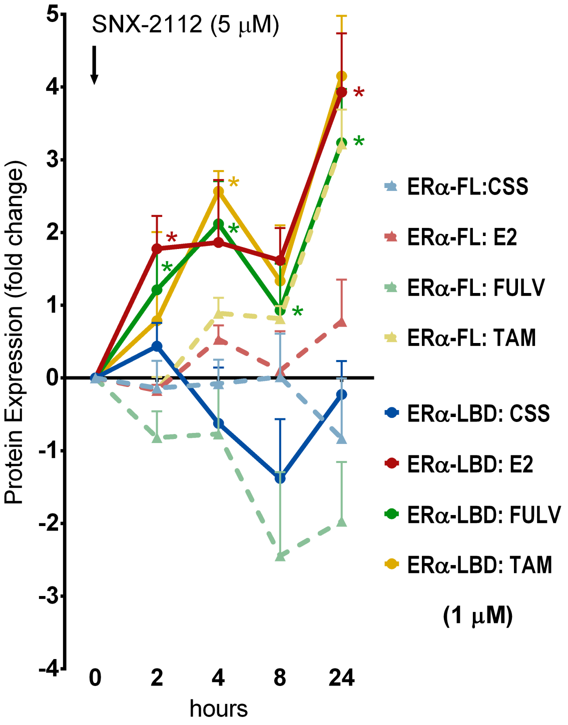

SNX-2112 (5 µM)
Protein Expression (fold change)
5
4
3
2
1
0
-1
-2
-3
-4
0
2
4
8
24
hours
ERα-FL:CSS
ERα-FL: E2
ERα-FL: FULV
ERα-FL: TAM
ERα-LBD: CSS
ERα-LBD: E2
ERα-LBD: FULV
ERα-LBD: TAM
(1 µM)

**Supplementary Fig. S1**

Fulvestrant resistant BC cell lines express a smaller ERα isoform, lacking the N-term domains and stabilized by ERα ligands. **A,** Proliferation of different breast cancer (BC) cell lines, in the presence or absence of fulvestrant 1 μM treatment, assayed using resazurin reagent and expressed as fluorescence intensity (adsorbance at 590 nm), taken as index of cell growth. Data are shown as mean ± s.e.m. ($n$ = 3 independent experiments). ** $P < 0.01$, two-way ANOVA. For each cell line, the two biological groups (vehicle vs. fulvestrant) were statistically compared and $P$ values are shown (two-way ANOVA, Sidak's test for multiple comparisons). FulvRes = fulvestrant resistant. **B,** Representative images of MCF-7 cells. Both WT and FulvRes cells were cultured in the presence or absence of fulvestrant 1 μM, for 7 days before image acquisition. A scale bar for each image representing 100 μm is shown. Magnification: 20X. **C,** Protein lysates from MCF-7 FulvRes and MDA-MB-231 (both treated with fulvestrant 1 μM for 24h) were immunoprecipitated using anti-ERα antibody. IP samples were run on two electrophoresis gels, one used for staining and band extraction, the other for WB check. Extracted bands were analyzed by capLC-MS/MS and results are shown. Sequenced/identified peptides from ERα protein are highlighted in dark grey and compared to the full-length ERα AA sequence. Different ERα domains with specific color code are also shown. AF1: transcription Activation Function-1 (cyan); DBD: DNA Binding Domain (red); Hinge (green); LBD/AF2: Ligand Binding Domain and transcription Activation Function-2 (yellow); amino acids involved in E2 binding (magenta); ERα-36 unique C-terminal (purple). **D,** Analysis on ERα-LBD sequence. **E,** Western blot analysis of ERα protein expression in breast BC cell lines. Cells were cultured for 24 h in the presence of vehicle (-) or fulvestrant 1 μM (+) before lysis. Vehicle = DMSO. ERα protein expression was normalized against β-actin. Increasing levels of hormonal therapy resistance (HTR) are indicated. **F,** ERα protein stability assay. ERα-FL and ERα-LBD protein levels were analyzed by western blot in MCF-7 and MDA-MB-231 cells, respectively. Protein samples were collected at different time points (2-24 h), after treatment. ERα protein levels were quantified and plotted as fold-change (log2), relative to SNX-2112 treatment alone (lane #2). CSS = charcoal stripped serum. Data are presented as mean ± s.e.m. ($n$ = 3 independent experiments); * $P < 0.01$, two-way ANOVA (Tukey's correction), ERα-LBD vs. ERα-FL.

### Supplementary Fig. S2

A

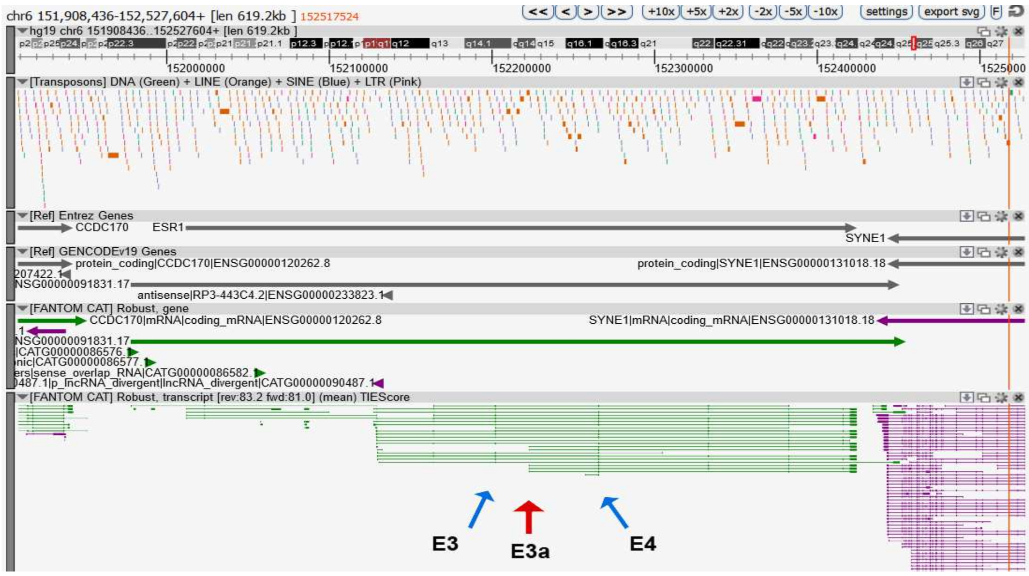


B

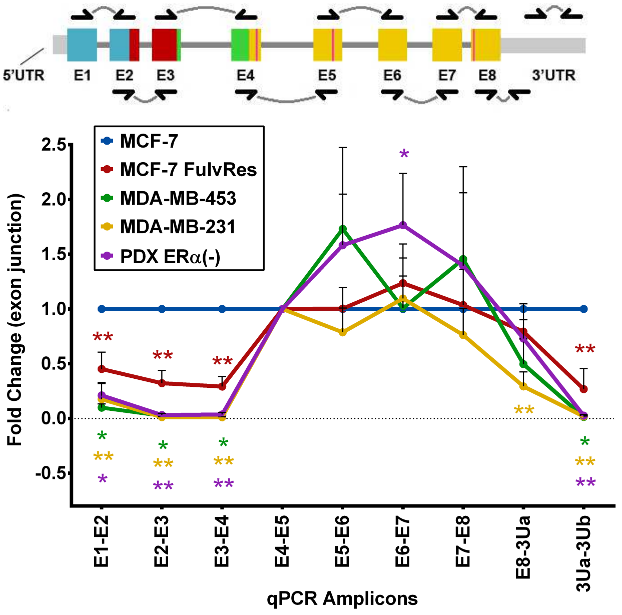


C

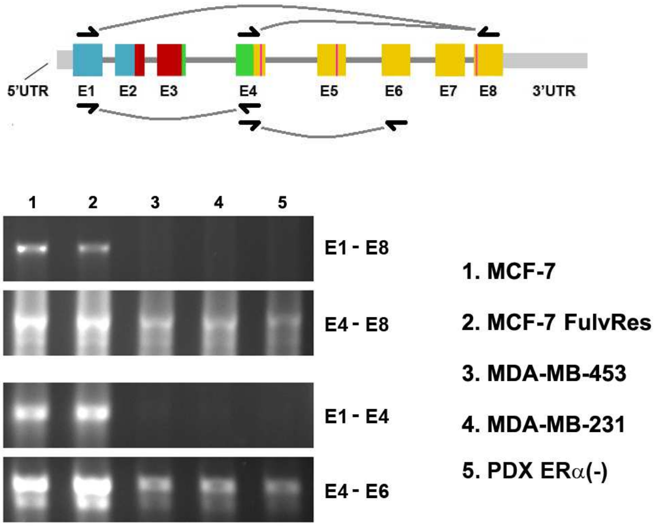


D

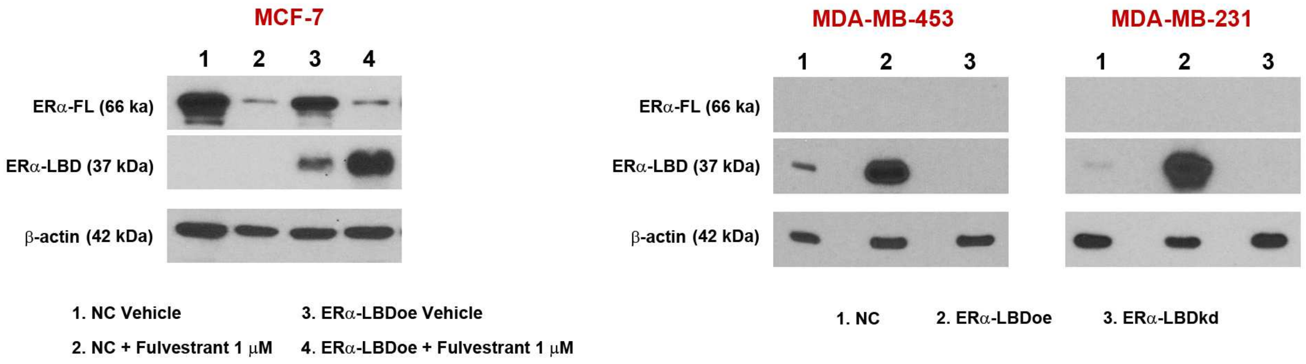

E

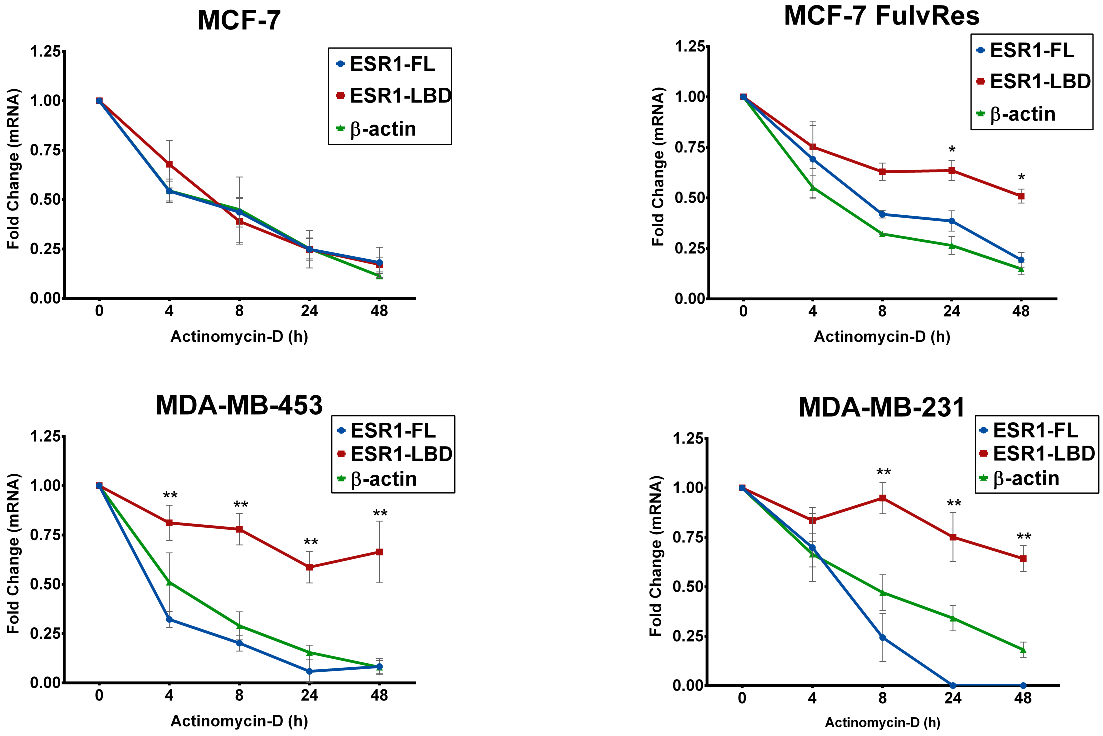
MCF-7
ESR1-FL
ESR1-LBD
β-actin
Fold Change (mRNA)
1.25
1.00
0.75
0.50
0.25
0.00
0
4
8
24
48
Actinomycin-D (h)
MCF-7 FulvRes
*
*
MDA-MB-453
**
MDA-MB-231
**

F

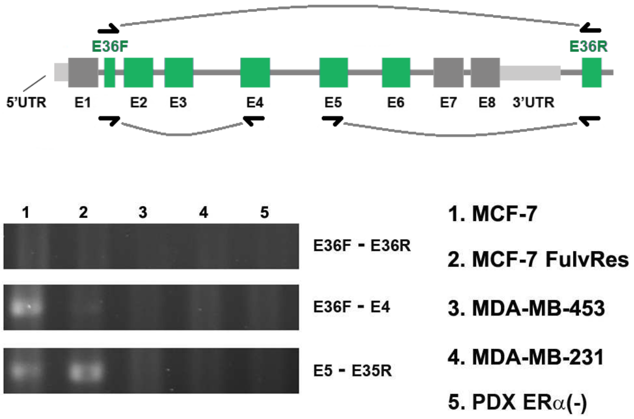
E36F
E36R
5'UTR
E1
E2
E3
E4
E5
E6
E7
E8
3'UTR
1
2
3
4
5
E36F - E36R
E36F - E4
E5 - E35R
1. MCF-7
2. MCF-7 FulvRes
3. MDA-MB-453
4. MDA-MB-231
5. PDX ERα(-)

G

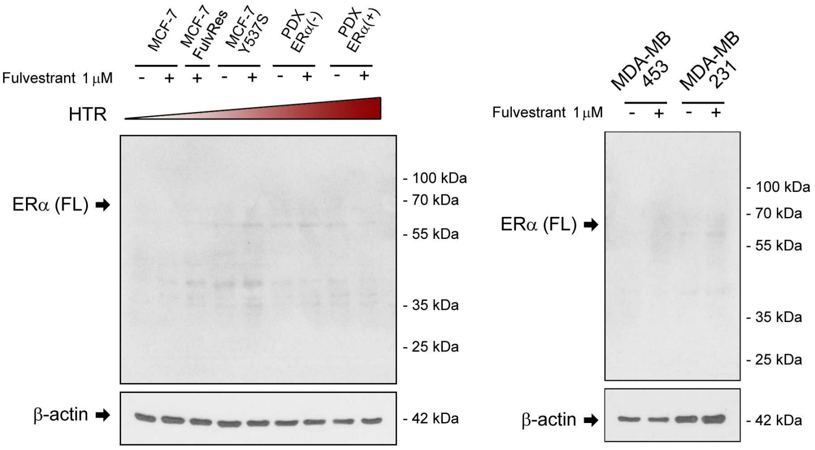
MCF-7
MCF-7 FulvRes
MCF-7 Y537S
PDX ERα(-)
PDX ERα(+)
Fulvestrant 1 μM
- + - + - + - + - +
HTR
ERα (FL)
β-actin
- 100 kDa
- 70 kDa
- 55 kDa
- 35 kDa
- 25 kDa
- 42 kDa
MDA-MB 453
MDA-MB 231
- + - +

H

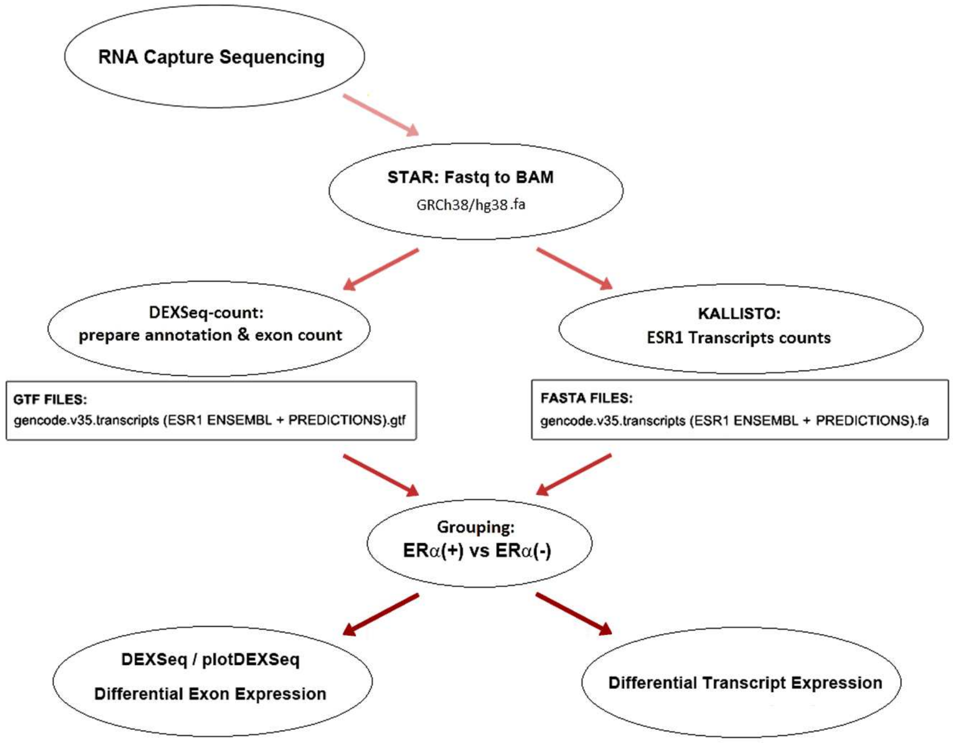
RNA Capture Sequencing
STAR: Fastq to BAM
GRCh38/hg38.fa
DEXSeq-count:
prepare annotation & exon count
KALLISTO:
ESR1 Transcripts counts
GTF FILES:
gencode.v35.transcripts (ESR1 ENSEMBL + PREDICTIONS).gtf
FASTA FILES:
gencode.v35.transcripts (ESR1 ENSEMBL + PREDICTIONS).fa
Grouping:
ERα(+) vs ERα(-)
DEXSeq / plotDEXSeq
Differential Exon Expression
Differential Transcript Expression

**Supplementary Fig. S2**

ERα-LBD is encoded by an *ESR1* transcript variant, having an alternative 5'UTR and characterized by the shortening of the 3'UTR sequence. ESR1-LBD transcript has higher stability, compared to ESR1-FL. ERα-LBD differs from ERα-36 isoform, at both transcript and protein level. **A,** Screenshot of ZEMBU Genome Browser showing data from FANTOM5/FANTOM CAT analyses collected on ESR1 gene. Exon 3 (E3) and exon 4 (E4) of ESR1 gene are highlighted by blue arrows, together with a new putative transcription starting site (TSS) mapped in the intronic region between the two exons (E3a, red arrow). **B,** Expression of ESR1 mRNA exon junctions in BC cell lines, analyzed by qPCR. Plot shows expression fold change, normalized on RPLP0 and relative to MCF-7 sample. Color-code of exons is based on different ERα protein domain. Data are presented as mean ± s.e.m. ($n$ = 3 independent experiments). * $P < 0.05$, ** $P < 0.01$, two-way ANOVA (Fisher's LSD test). **C,** Expression of different ESR1 mRNA regions in BC cell lines, analyzed by RT-PCR. The scheme above depicts PCR amplicons and their position on ESR1 transcript. Color-code of exon as in (**B**). **D,** Western blot analysis of ERα protein variants expression (ERα-FL vs. ERα-LBD) in different BC cell lines (stable clones). Vehicle (DMSO 0.01%) or fulvestrant treatment (1 µM, 24 h) was added to MCF-7 cells (on the left). NC: cells transduced with control vector. ERα-LBDoe: cells transduced with ESR1-LBD CDS sequence (exon 4 to exon 8), inducing ERα-LBD protein overexpression. ERα-LBDkd: cells transduced with CRISPR/CAS9 vector targeting ESR1 exon 4 and promoting ERα-LBD protein knockdown. **E,** BC cells lines were treated with actinomycin-D and tested by qPCR at different time points for ESR1 and β-actin mRNA expression. ESR1-FL: PCR amplicon ranging from ESR1 exon E1 to E3. ESR1-LBD: PCR amplicon ranging from ESR1 exon E4 to E5. β-actin transcript was taken as positive control for mRNA decay. Plots shows mRNA fold change, normalized on RPLP0 and relative to time 0. Data are presented as mean ± s.e.m. ($n$ = 3 independent experiments). * $P < 0.05$, ** $P < 0.01$, two-way ANOVA (Fisher's LSD test). **F,** Expression of different ESR1 mRNA regions in BC cell lines, analyzed by RT-PCR. The scheme above depicts PCR amplicons and their position on ESR1 transcript. Exons of ERα-36 transcript are in green. **G,** Western blot analysis of ERα-36 protein expression in BC cell lines. Cells were cultured for 24 h in the presence of vehicle (-) or fulvestrant 1 µM (+) before lysis. Vehicle = DMSO. ERα protein expression was normalized against β-actin. Increasing levels of hormonal therapy resistance (HTR) are indicated. **H,** A schematic summary of the pipeline used to analyze RNA sequencing raw data obtained from RNA capture-seq experiment.

### Supplementary Fig. S3

**A**

| *Anti-ERa (Cell Signaling)* | MCF-7 | | MCF-7 +Fulv | | MCF-7 FulvRes +Fulv | |
|---|---|---|---|---|---|---|
| | Nucl | Mito | Nucl | Mito | Nucl | Mito |
| threshold A | 50 | 50 | 50 | 50 | 50 | 50 |
| threshold B | 50 | 50 | 50 | 50 | 50 | 50 |
| number of colocalized voxels | 535701 | 28128 | 75341 | 39987 | 49694 | 96212 |
| % of dataset colocalized | 1.7 | 0.09 | 0.27 | 0.14 | 0.23 | 0.44 |
| % of ROI colocalized | 1.7 | 0.09 | 0.27 | 0.14 | 0.23 | 0.44 |
| % of volume A above threshold colocalized (DAPI / OXPHOS) | 34.71 | 4.4 | 29.1 | 10.63 | 30.14 | 46.75 |
| **% of volume B above threshold colocalized (ERa)** | **53.89** | **2.83** | **31.28** | **16.6** | **13.55** | **26.24** |
| % of material A above threshold colocalized (DAPI / OXPHOS) | 35.48 | 4.91 | 29.52 | 11.89 | 29.24 | 50.07 |
| **% of material B above threshold colocalized (ERa)** | **56.09** | **2.06** | **34.49** | **14.63** | **14.94** | **25.42** |
| % of ROI material A colocalized (DAPI / OXPHOS) | 10.77 | 2.85 | 7.46 | 8.42 | 7.66 | 37.34 |
| **% of ROI material B colocalized (ERa)** | **31.29** | **1.15** | **15.58** | **6.61** | **9.89** | **16.82** |
| Pearson's coefficient in dataset volume | 0.5687 | 0.0975 | 0.5852 | 0.3261 | 0.452 | 0.4067 |
| Pearson's coefficient in ROI volume | 0.5687 | 0.0975 | 0.5852 | 0.3261 | 0.452 | 0.4067 |
| Pearson's coefficient in colocalized volume | -0.0016 | -0.1769 | 0.0433 | -0.0152 | -0.0603 | 0.0343 |
| original Mander's coefficient A | 0.7252 | 0.8992 | 0.7513 | 0.9647 | 0.6997 | 0.9805 |
| original Mander's coefficient B | 0.996 | 0.4609 | 0.9714 | 0.6228 | 0.9063 | 0.5377 |
| thresholded Mander's coefficient A | 0.1586 | 0.0551 | 0.145 | 0.1061 | 0.2041 | 0.4425 |
| thresholded Mander's coefficient B | 0.4157 | 0.0532 | 0.2268 | 0.1976 | 0.132 | 0.2136 |

| *Anti-ERa (Cell Signaling)* | MDA-MB-453 | | MDA-MB-231 | | PDX ERα(-) | |
|---|---|---|---|---|---|---|
| | Nucl | Mito | Nucl | Mito | Nucl | Mito |
| threshold A | 50 | 50 | 50 | 50 | 50 | 50 |
| threshold B | 50 | 50 | 50 | 50 | 50 | 50 |
| number of colocalized voxels | 2634 | 90200 | 23806 | 81216 | 4412 | 78345 |
| % of dataset colocalized | 0.01 | 0.3 | 0.11 | 0.37 | 0.02 | 0.31 |
| % of ROI colocalized | 0.01 | 0.3 | 0.11 | 0.37 | 0.02 | 0.31 |
| % of volume A above threshold colocalized (DAPI / OXPHOS) | 3.77 | 33.87 | 6.65 | 45.95 | 2.79 | 31.79 |
| **% of volume B above threshold colocalized (ERa)** | **0.67** | **22.8** | **10.9** | **37.19** | **2.16** | **38.28** |
| % of material A above threshold colocalized (DAPI / OXPHOS) | 3.77 | 39.11 | 6.43 | 53.84 | 2.73 | 35.49 |
| **% of material B above threshold colocalized (ERa)** | **0.53** | **22.47** | **9.27** | **41.03** | **1.8** | **38.27** |
| % of ROI material A colocalized (DAPI / OXPHOS) | 0.31 | 23.7 | 4.55 | 40.54 | 0.61 | 23.12 |
| **% of ROI material B colocalized (ERa)** | **0.26** | **11.13** | **3.66** | **16.19** | **0.79** | **16.89** |
| Pearson's coefficient in dataset volume | 0.2851 | 0.4097 | 0.3069 | 0.5341 | 0.3525 | 0.5341 |
| Pearson's coefficient in ROI volume | 0.2851 | 0.4097 | 0.3069 | 0.5341 | 0.3525 | 0.5341 |
| Pearson's coefficient in colocalized volume | -0.0453 | 0.0104 | -0.0515 | 0.1891 | -0.0215 | -0.0183 |
| original Mander's coefficient A | 0.7753 | 0.9737 | 0.9518 | 0.9927 | 0.7565 | 0.9925 |
| original Mander's coefficient B | 0.9236 | 0.6657 | 0.7661 | 0.5258 | 0.9522 | 0.8155 |
| thresholded Mander's coefficient A | 0.0735 | 0.3069 | 0.0774 | 0.4472 | 0.0621 | 0.2852 |
| thresholded Mander's coefficient B | 0.0127 | 0.1644 | 0.1685 | 0.2169 | 0.0576 | 0.2711 |

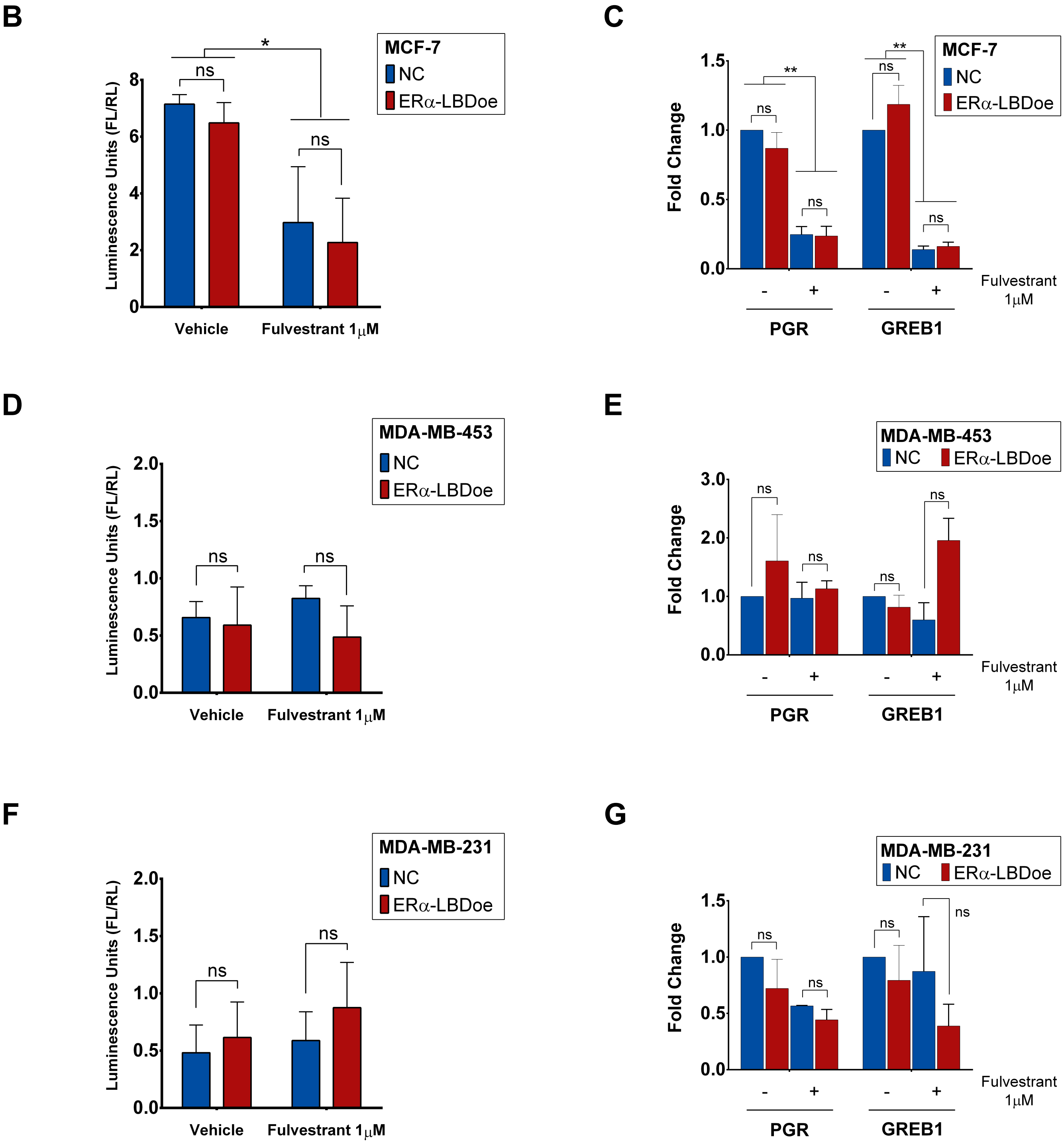


**Supplementary Fig. S3**

Confocal images analysis and statistics indicate cytoplasmic and mitochondrial localization for ERα-LBD in breast cancer cells. ERα-LBD does not activate transcription in the nucleus. **A,** Table summarizing statistics from confocal analysis on different BC cell lines. In particular, colocalization data are shown. Blue and red numbers indicate colocalization level of ERα protein into the nucleus or mitochondria of cells, respectively. **B, D** and **F,** ERE (Estrogen Responsive Element) promoter-driven luciferase reporter assay in MCF-7, MDA-MB-453 and MDA-MB-231 cells, respectively. Firefly luciferase levels were normalized to Renilla luciferase. Controls (NC) were compared to cells overexpressing ERα-LBD (oe), either in the presence or absence of fulvestrant 1 μM treatment (24 h). Vehicle = DMSO 0.01%. Data are presented as mean ± s.e.m. ($n$ = 3 independent experiments). * $P < 0.05$, ns = not significant; two-way ANOVA (Sidak's correction). **C, E** and **G,** Expression of PGR and GREB1 mRNA in MCF-7, MDA-MB-453 and MDA-MB-231 cells, respectively. Analysis was carried out by qPCR. Samples and treatments: same as above. Plot shows fold change expression, relative to NC without treatment. In all panels, blue indicates NC samples and red indicates ERα-LBDoe/kd samples. Data are presented as mean ± s.e.m. ($n$ = 6 or $n$ = 4 independent experiments). ** $P < 0.01$, ns = not significant; two-way ANOVA (Sidak's correction).

### Supplementary Fig. S4

**A**

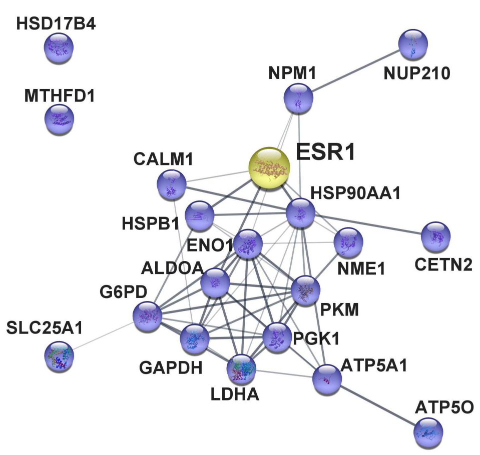


**Glycolysis and Gluconeogenesis**

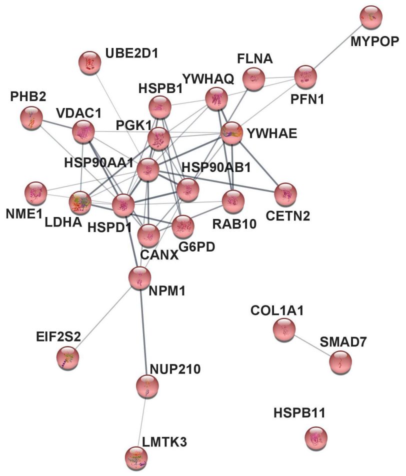


**Signaling**

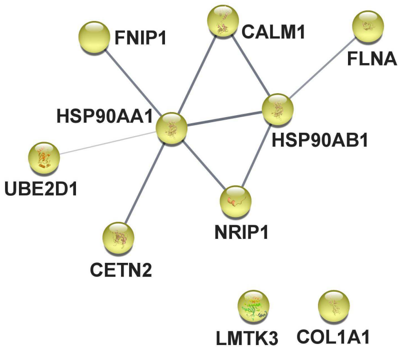


**Hypoxia and Angiogenesis**

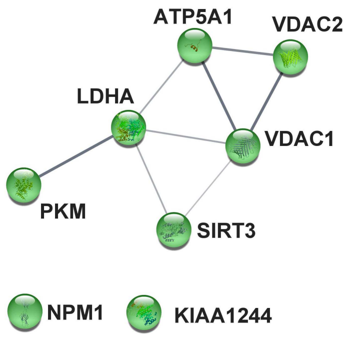


**Mitochondrial Metabolism and Respiration**

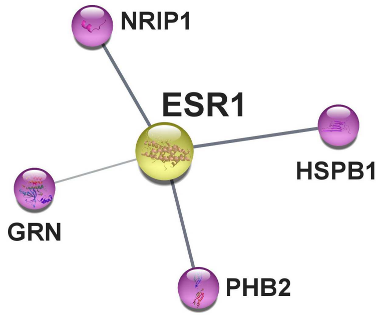


**ERα Signaling**

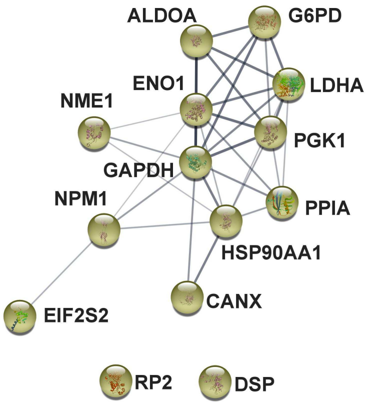


**MYC Signaling**

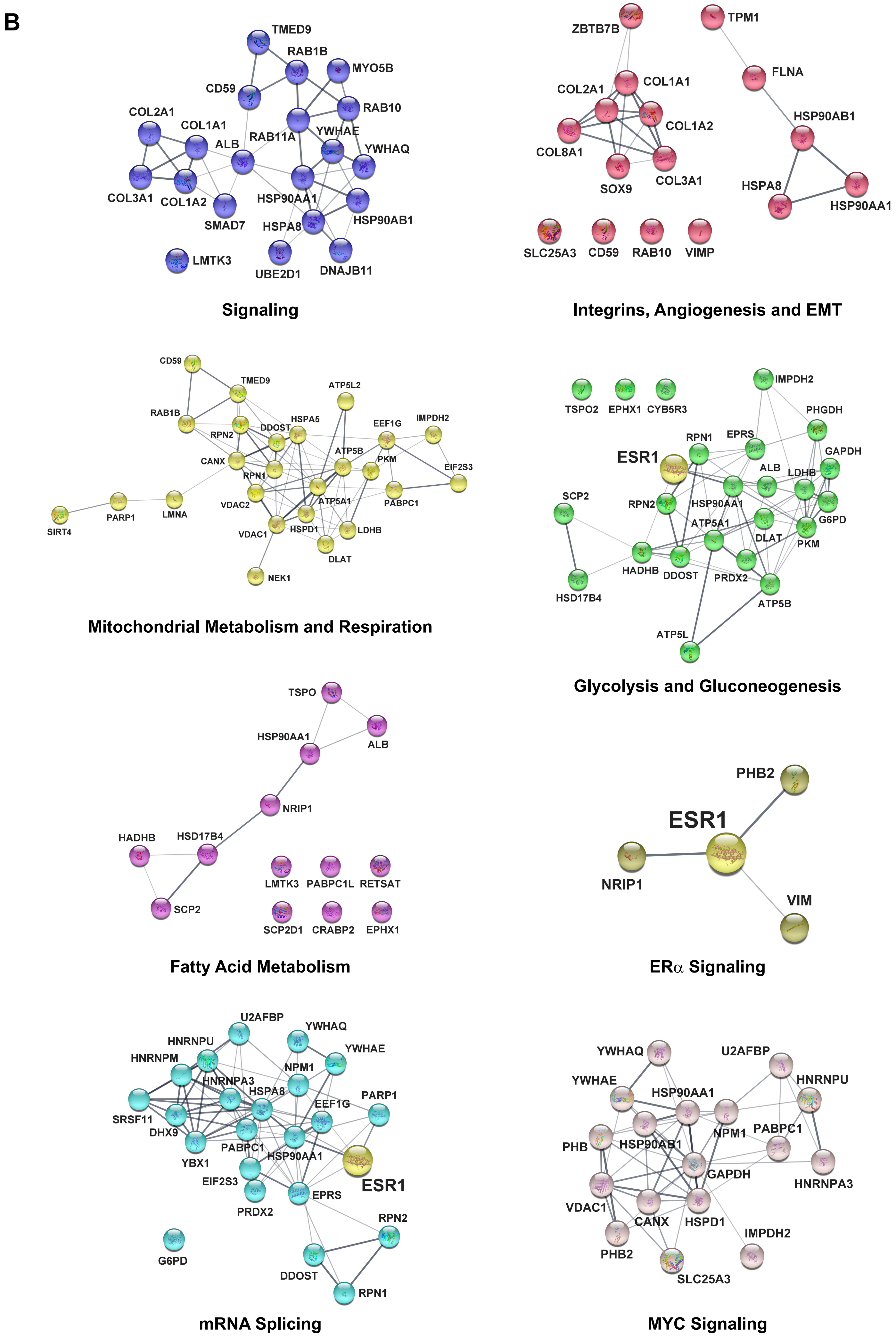
B
TMED9
RAB1B
MYO5B
CD59
RAB10
COL2A1
COL1A1
ALB
RAB11A
YWHAE
YWHAQ
COL3A1
COL1A2
SMAD7
HSP90AA1
HSPA8
HSP90AB1
LMTK3
UBE2D1
DNAJB11
Signaling
ZBTB7B
TPM1
FLNA
COL2A1
COL1A1
COL1A2
COL8A1
SOX9
COL3A1
HSP90AB1
HSPA8
HSP90AA1
SLC25A3
CD59
RAB10
VIMP
Integrins, Angiogenesis and EMT
CD59
TMED9
ATP5L2
RAB1B
HSPA5
EEF1G
IMPDH2
RPN2
DDOST
ATP5B
CANX
PKM
EIF2S3
RPN1
ATP5A1
PABPC1
VDAC2
SIRT4
PARP1
LMNA
VDAC1
HSPD1
LDHB
DLAT
NEK1
Mitochondrial Metabolism and Respiration
TSPO2
EPHX1
CYB5R3
IMPDH2
PHGDH
RPN1
EPRS
ESR1
GAPDH
ALB
LDHB
SCP2
RPN2
HSP90AA1
G6PD
ATP5A1
DLAT
PKM
HADHB
DDOST
PRDX2
HSD17B4
ATP5B
ATP5L
Glycolysis and Gluconeogenesis
TSPO
HSP90AA1
ALB
NRIP1
HADHB
HSD17B4
SCP2
LMTK3
PABPC1L
RETSAT
SCP2D1
CRABP2
EPHX1
Fatty Acid Metabolism
PHB2
ESR1
NRIP1
VIM
ERα Signaling
U2AFBP
YWHAQ
HNRNPU
YWHAE
HNRNPM
NPM1
HNRNPA3
HSPA8
PARP1
EEF1G
SRSF11
DHX9
PABPC1
HSP90AA1
YBX1
EIF2S3
ESR1
EPRS
PRDX2
RPN2
G6PD
DDOST
RPN1
mRNA Splicing
YWHAQ
U2AFBP
YWHAE
HNRNPU
HSP90AA1
PHB
NPM1
PABPC1
HSP90AB1
GAPDH
HNRNPA3
VDAC1
CANX
HSPD1
IMPDH2
PHB2
SLC25A3
MYC Signaling

**Supplementary Fig. S4**

ERα-LBD interacts with several proteins from different cell compartments. **A** and **B,** Analysis of ERα-LBD protein-protein interaction (PPI) network in MCF-7 and TNBC models respectively, using Cytoscape and STRING software. Proteins are presented as nodes. PPIs networks associated to specific biological functions are presented by using different colors.

### Supplementary Fig. S5

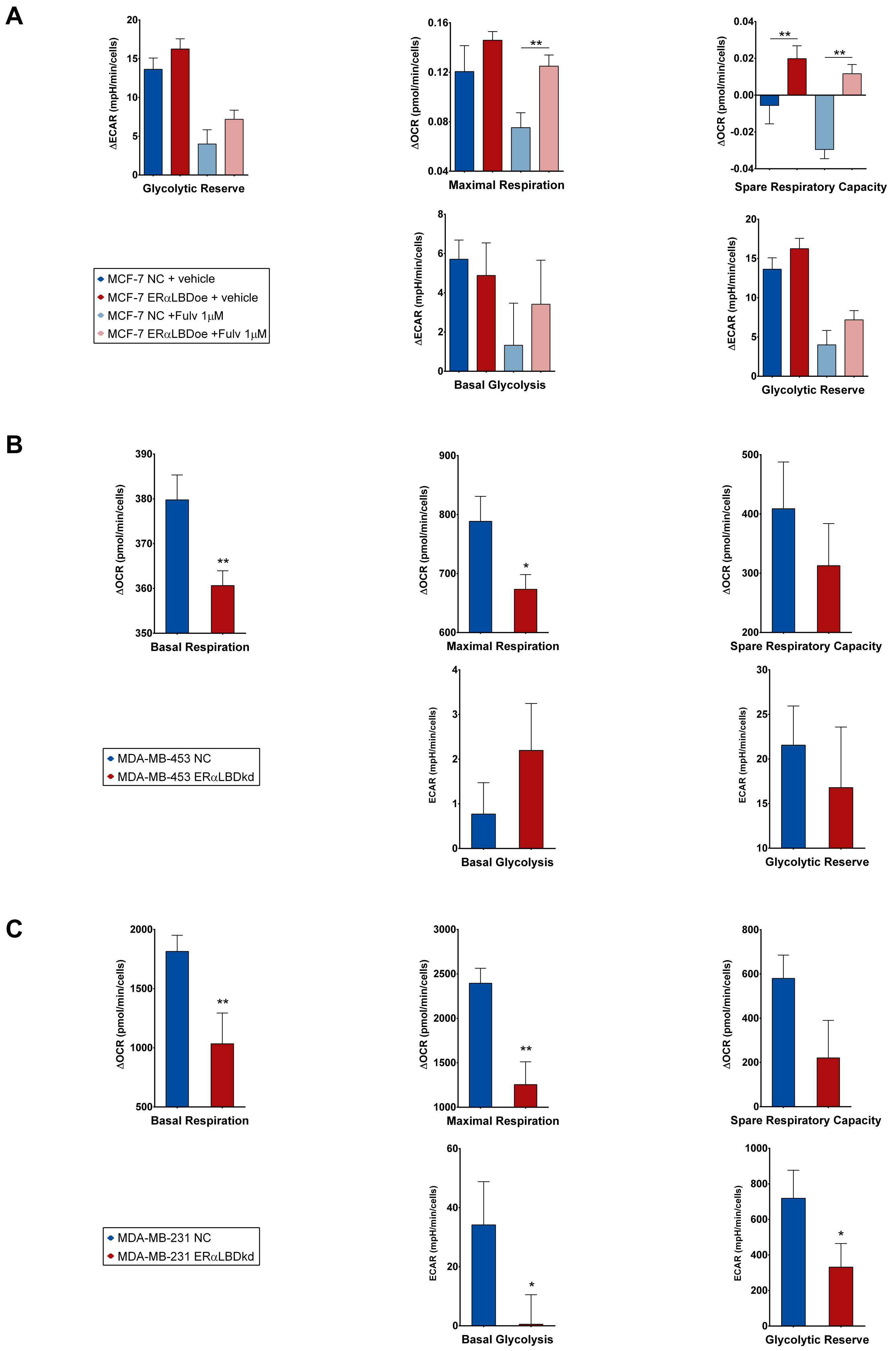

**Supplementary Fig. S5**

ERα-LBD levels affect respiratory and glycolytic parameters in BC cells. **A,** Evaluation of respiratory and glycolytic parameters in MCF-7 cell clones. Fulvestrant 1 µM (24 h pre-treatment) or vehicle (DMSO) was added to cells. ERα-LBD overexpressing (oe) cells were compared to controls (NC). Analyses were carried out by using XF Cell Mito Stress Kit (Agilent). Following manufacturer's guidelines, the calculation of all parameters was based on ΔOCR and ΔECAR values collected during the assay. **B** and **C,** Evaluation of respiratory and glycolytic parameters in TNBC cells (MDA-MB-453/-231). Cells with ERα-LBD knockdown (kd) were compared to controls (NC). Analysis was carried out as described above. Data in the figures are presented as mean ± s.e.m. ($n$ = 2 independent experiments). * $P < 0.05$, ** $P < 0.01$; unpaired t test, one-sided.

### Supplementary Fig. S6

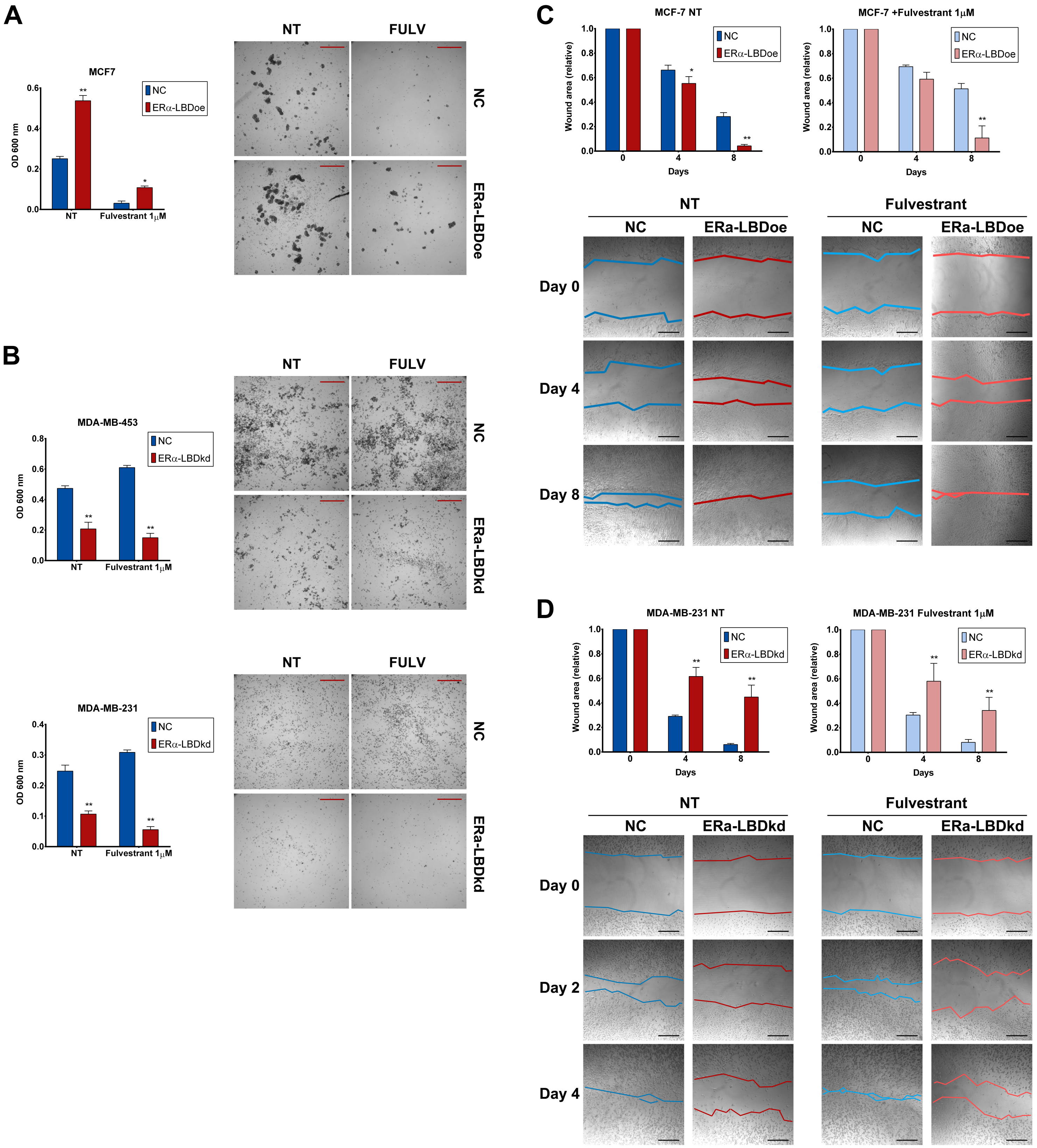


**Supplementary Fig. S6**

ERα-LBD levels affect BC cells 3-D growth and migration. **A,** 3-D growth of BC stable clones, measured as optical density (OD) at 600 nm of cell suspensions. ERα-LBD overexpression (MCF-) or knockdown (MDA-MB-453 & 231) was compared to controls (NC), either in the absence (NT) or presence of fulvestrant 1 µM treatment. Representative images of cell 3-D growth are also shown. Fulv = fulvestrant 1 µM. Scale bar: 200 µm. Magnification: 10X. **B** and **C,** Cell migration of BC stable clones was tested by wound healing assay, at different time points (days). Samples and treatments: same as in (**A**). In all panels, blue indicates NC samples and red indicates ERαLBDoe/kd samples. Representative images of scratches are also shown, with wound edges highlighted by different colored lines. Fulv = fulvestrant 1 µM. Scale bar: 200 µm. Magnification: 10X. All data in the figure are presented as mean ± s.e.m. (*n* = 3 independent experiments), * $P < 0.05$, ** $P < 0.01$, two-way ANOVA (Sidak's correction).

### Supplementary Fig. S7

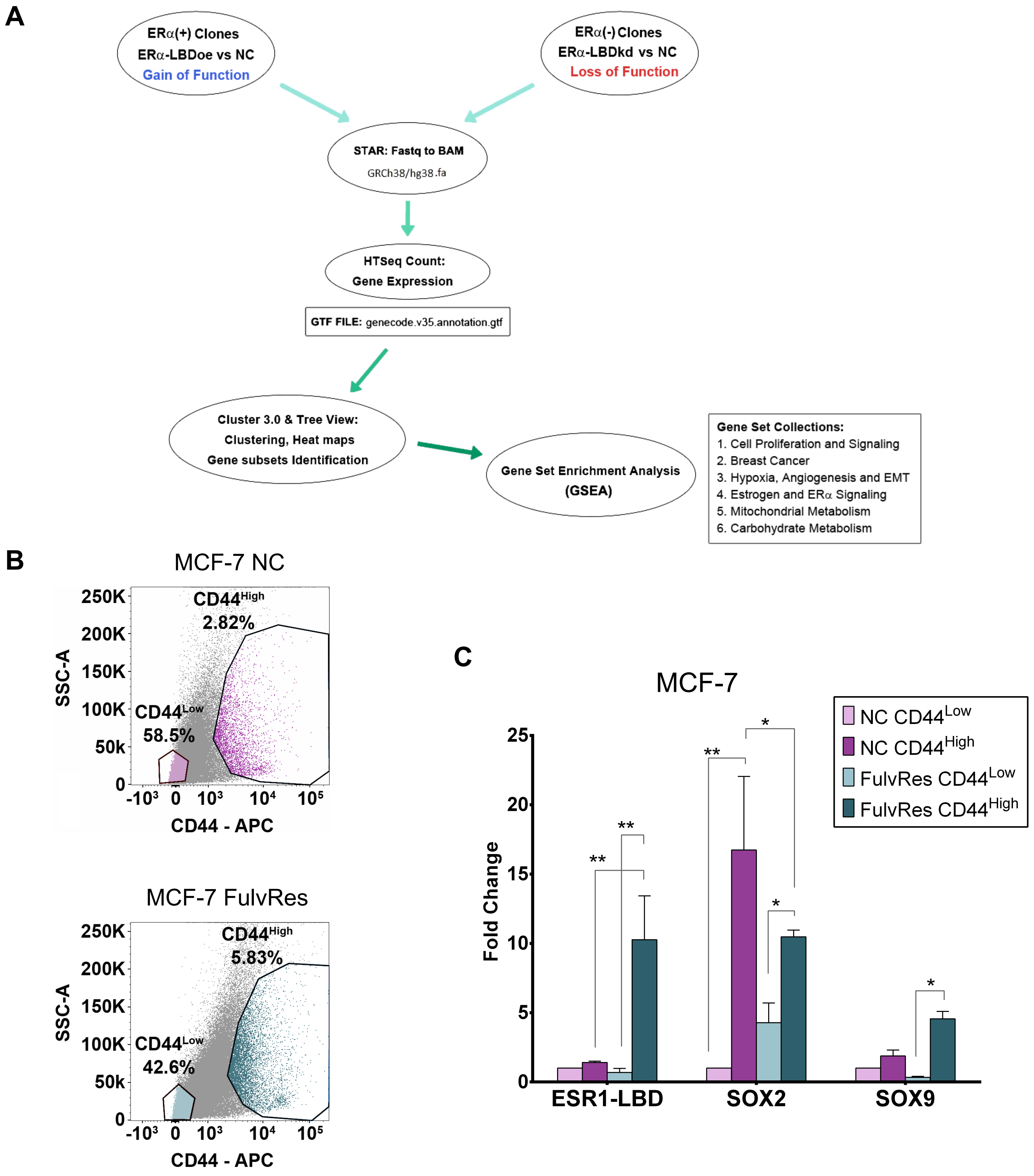


**Supplementary Fig. S7**

RNA-seq analysis of ERα-LBD 'gain-/ loss-of-function' and fulvestrant resistant BC models. ERα-LBD expression in stem-like cells. **A,** Schematic summary of the pipeline used to analyze RNA-seq raw data from experiments on BC cell clones. **B,** Representative images of flow cytometry analysis of CD44$^{Low}$ and CD44$^{High}$ cells, isolated from BC murine xenografts models (MCF-7 and MCF-7 FulvRes). Color code: purple, MCF-7 NC; teal, MCF-7 FulvRes; light, CD44$^{Low}$; dark, CD44$^{High}$. **C,** Total RNA was extracted from CD44$^{Low/High}$ sorted BC cells and analyzed by qPCR. Plot shows levels of ESR1-LBD (normalized first on RPLP0, then on ESR1-FL) and stem-cell markers Sox2 and Sox9 (normalized on RPLP0), expressed as fold change (CD44$^{High}$ vs. CD44$^{Low}$). Color code as in (**B**). Data are presented as mean ± s.e.m. ($n$ = 2 independent experiments; $n$ = 3 replicates, each experiment). * $P < 0.05$, ** $P < 0.01$, two-way ANOVA (Fisher's LSD test) and unpaired t test, two-tailed.

### Supplementary Fig. S8

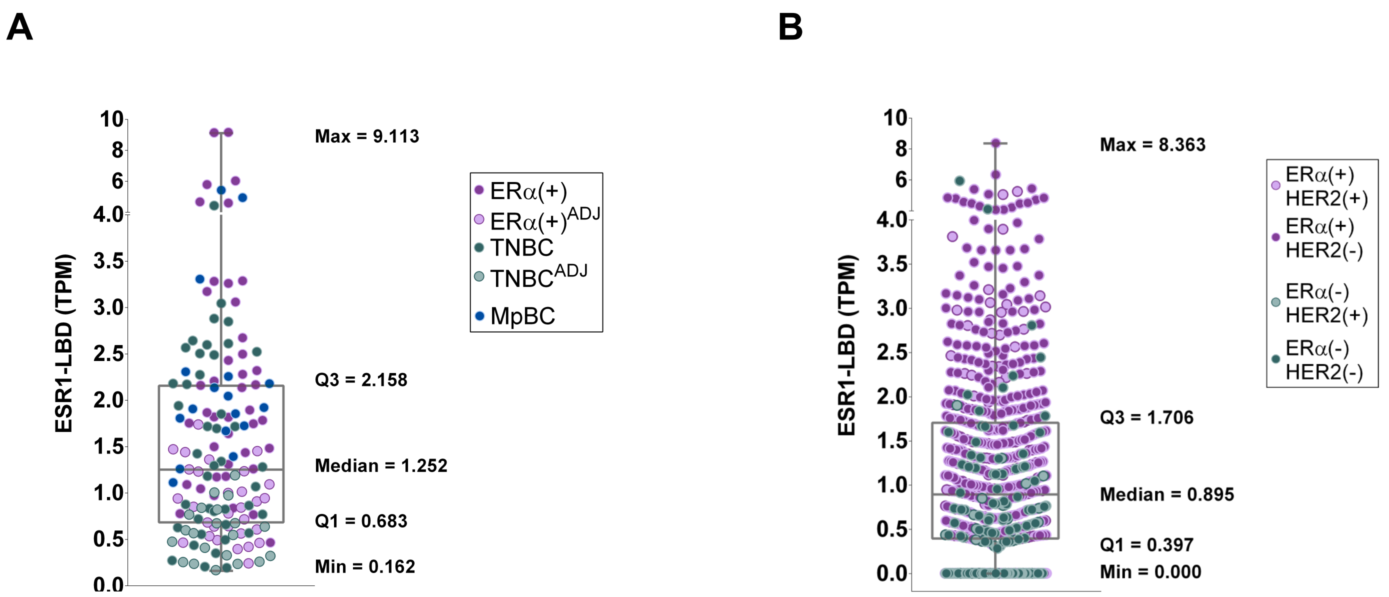


**Supplementary Fig. S8**

Analysis of ESR1-LBD expression in tumor samples from BC patients, with statistics. **A** and **B,** Box and whiskers plots representing the distribution of ESR1-LBD expression values (TPM) from all BC or TCGA-BRCA samples, respectively. Median, Q1/Q3 and Min/Max values are shown. Please refer to the main text for the description and color code of samples/groups (**Fig. 8A** and **E**).
